# Granulin^+^ macrophages promote lineage plasticity in prostate cancer through paracrine signaling loops

**DOI:** 10.1101/2025.07.09.663835

**Authors:** Zhipeng Zhu, Mi Zhang, Ying Song, Wei Jiang, Fang Cao, Yicong Yao, Xiaotong Yu, Hongyu Zhao, Husile Baiyin, De Chang, Denglong Wu, Xiaolu Zhao, Gang Wu, Kailong Li, Fengbiao Mao

## Abstract

The role of tumor-infiltrating immune cells in driving phenotype switching remains unclear, despite the well-established association between lineage transition and drug resistance in prostate cancer. This study employed an integrated analysis of single-cell multiomics to investigate the dynamics of immune infiltration, transcriptional programs, and cell-cell communication in prostate cancer. Our results demonstrated that granulin (GRN) positive macrophages facilitated the transition from adenocarcinoma to a multilineage state with mesenchymal and stem-like traits by activating intra-tumoral NF-κB signaling. Subsequently, the multilineage clones induced macrophages to highly express granulin through the secretion of CSF1, forming a positive feedback cell communication loop. Next, we validated the biological function of granulin in mediating epithelial-mesenchymal transition in vitro. Additionally, organoids drug resistance assay demonstrated that granulin drove resistance to androgen receptor (AR)-targeted drug (enzalutamide). Moreover, pharmacologic blockade of the CSF-1/CSF-1R axis in TRAMP mouse models reduced the expression of GRN in macrophages and suppressed the formation of multilineage subclones in prostate malignant cells. Furthermore, multiplex immunofluorescence staining of tumor samples from TRAMP mouse models revealed the VIM lineages were spatially in close contact with macrophages. Meanwhile, Cytometry by Time-Of-Flight (CyTOF) analysis validated our findings at single-cell protein level in patients with castration-resistant prostate cancer. Besides, three distinct tumor-infiltrating subsets associated with disease relapse were identified, including DCN^+^ endothelial cells, CCL7^+^ fibroblasts, and IFIT1^+^ neutrophils. These results offer potential therapeutic targets to address lineage plasticity-driven resistance to AR-targeted therapy.

## Introduction

Prostate cancer is a prevalent malignancy among males globally, characterized by its clinical heterogeneity[1]. Aggressive forms of the disease, particularly in patients with locally advanced or metastatic tumors, frequently develop rapid resistance to the commonly used androgen deprivation therapy (ADT), leading to the emergence of castration-resistant prostate adenocarcinoma (CRPC-Adeno), which is characterized by high androgen receptor (AR) signaling in the luminal lineage[2]. The re-establishment of the AR-driven transcriptional program or the activation of alternative transcription factors to circumvent AR signaling may contribute to the resistance to ADT[3]. Newer drugs, such as enzalutamide and abiraterone, are directed toward suppressing AR signaling to prolong survival but seem to induce various phenotypes independent to AR signaling, including mixed prostatic adenocarcinomas with neuroendocrine (NE) features, small-cell carcinomas, or double-negative features[4–9]. Lineage plasticity is a novel mechanism that drives multiple lineage differentiation to confer therapeutic resistance independent of AR activity[10]. Studies generally agreed that lineage plasticity contributes to the transition of luminal adenocarcinoma cells to stem cell-like, epithelial-mesenchymal transition (EMT)-like, and multilineage states that may redifferentiate into various lineages[11, 12], such as the NE lineage. Although genetic alterations[13–15] and epigenetic alterations[15–19] are recognized as candidate regulators of lineage plasticity in prostate cancer, druggable targets responsible for prostate cancer lineage plasticity remain elusive.

Conversely, there is an increasing consensus indicating that non-genetic modulation can determine cancer cell state to shape metastatic and drug-resistant features, with the main signal sources for non-genetic modulation originating from the tumor microenvironment (TME). For instance, SPP1 from cancer-associated fibroblast promotes castration resistance in prostate cancer[20], and KIT ligand from stromal cells drives lineage plasticity in prostate cancer[21]. The TME is enriched with numerous complex inflammatory factors that can synergistically interact with oncogenic mutations to drive hyperplasic and malignant transformation[12, 20, 22–24]. Macrophages are the primary sources of numerous inflammatory cytokines in prostate inflammation[25], particularly associated with poorly differentiated and plastic cells. For instance, the THP-1 human macrophage-like cell line has been shown to promote prostate hyperplasia[26], and macrophage cell lines secrete interleukin-6 (IL-6) to support neuroendocrine prostate cancer (NEPC) development[27]. However, further research is needed to explore the inflammatory factors released by macrophages that are indispensable for the maintenance of stem-like traits and lineage potential.

In this study, we performed single-cell multiomics analysis to explore the complexity of the TME and understand the underlying molecular processes involved in prostate cancer lineage plasticity. In addition, we validated the findings using cell co-culture in vitro, target intervention in vivo, organoids drug resistance assay and spatiotemporal information within the TME. Our study revealed that monocytes/macrophages promote the transition from adenocarcinoma to VIM lineage with mesenchymal and stem-like features through a granulin/NF-κB signaling paracrine mechanism. Conversely, the VIM lineages in turn facilitate monocytes/macrophages to highly express GRN via the secretion of CSF1, forming a positive feedback cell-cell communication loop. In summary, we found that paracrine feedback loop via granulin-NFκB-CSF1 axis may be a potential therapeutic target, and targeting this loop could potentially reverse AR signaling-targeted therapy resistance.

## Results

### Dynamic TME compositions contribute to cellular heterogeneity in prostate cancer

In a previous study, Chan et al. conducted single-cell RNA sequencing (scRNA-seq) on prostate cancer derived from GEMMs to investigate cellular heterogeneity and demonstrated lineage plasticity in prostate cancer depends on the cell-autonomous JAK/STAT inflammatory signaling[23]. However, it remains unclear which TME factors are crucial in driving the intermediate transcriptional cell state of malignant cells. To study the role of TME compositions in lineage plasticity, we re-analyzed scRNA-seq data from the 29 mice including 9 wildtype (WT), 7 Pten−/−Rb−/− (PtR) and 13 Pten−/−Rb−/−Trp53−/− (PtRP)], which reflect the progression from adenocarcinoma to NEPC (**Figure 1A**). We integrated 29 scRNA-seq datasets and manually annotated these single cells into 16 different cell lineages (Figure 1B, Figure S1). We investigated the cellular composition at different time points, specifically during the transition from adenocarcinoma to NEPC (Figure 1C). We found that Mono/Macro/DC exhibited significantly increasing in the early stage (PRP:8-9weeks_Intact) and then decreased in the later stage (PRP:12-16weeks_Intact) (Figure 1D). We further investigated the cancer cell differentiation trajectory and identified Adeno as the early lineage stage, TFF3, VIM and POU2F3 as intermediate lineage stages, but NEPC cell types as the late lineage stage through pseudotime analysis (Figure 1E). The tendency of lineage transition aligned with the timing of cancer evolution (Figure 1F), and was also in line with the alterations of cell composition throughout tumorigenesis (Figure 1G). These findings revealed that the composition of non-malignant and malignant cells exhibited a significant dynamics along with lineage transition.

**Figure 1:**
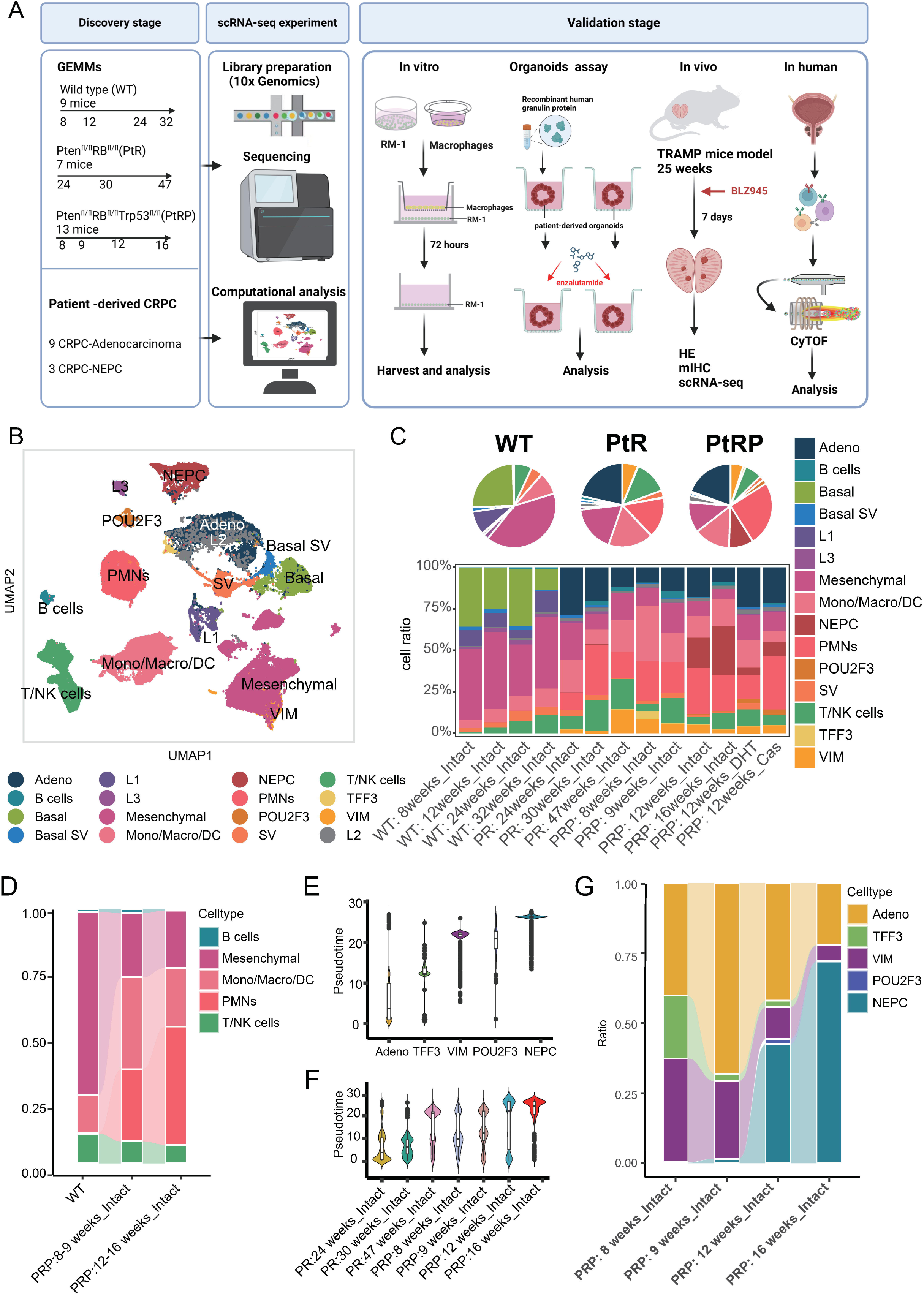
Single-cell multiomics analyses of 29 mice samples. (A) Schematic of this study design. Panel created in BioRender. Huafeng, F. (2025) https://BioRender.com/tg65p94 (**B**) UMAP visualization depicting cell populations colored by annotated major cell types. (**C**) Pie charts illustrating the distribution of cells colored by annotated major cell types assigned to sample types (above). Fraction of cells (y-axis) from samples at relevant timepoints colored by annotated major cell types (below). (**D**) Fraction of cells (y-axis) from samples at relevant timepoints colored by annotated tumor microenvironment cell types. (**E, F**) Violin plot showing pseudotime across five malignant lineages (**E**) and seven timepoints (**F**). (**G**) Fraction of cells (y-axis) from samples at relevant timepoints colored by annotated five malignant lineages.

### Tumor heterogeneity of prostate cancer lineages

We then investigated the tumor heterogeneity of malignant cells representing five distinct cell lineages. Unsupervised hierarchal clustering analysis revealed a distinct transcriptome pattern in the VIM lineage among five malignant lineages (**Figure 2A**). Meanwhile, the VIM lineage signature was associated with unfavorable patient survival based on RNA-seq data from TCGA (Figure 2B). To identify the activated molecular pathways in VIM lineage, we performed GSEA and revealed significant enrichment in inflammatory response, cytokine-receptor interactions and EMT signaling in VIM lineage (Figure 2D). EMT trait has been reported to endow cells with stem-like property, paving the way for redifferentiation into a new lineage[28]. We scored the cancer cells using EMT, stemness, and NF-κB signaling signature and observed VIM lineage exhibited mesenchymal and stem-like traits with excessive activation of NF-κB signaling (Figure S2A), coinciding with cancer progression stages (Figure S2B) and similar to previously reported drug-persistent mesenchymal stem-like PC (MSPC) cells[29]. The dynamic transdifferentiation model proposes that luminal adenocarcinoma cells undergo a transition to a partial EMT and stem-like state, followed by redifferentiation into a new lineage such as NE lineage[11]. Consistently, RNA velocity analysis showed a trajectory linking VIM lineage with Adeno and NEPC lineage (Figure 2C). Employing the cNMF algorithm, we further decomposed the malignant cells into nine consensus modules, each annotated with specific biological functions (Figure 2E). Importantly, EMT, biosynthesis, cell cycle, and NE2 corresponding to cancer lineage states, were projected on a two-dimensional branch. Intriguingly, the Adeno and TFF3 lineages were primarily enriched in the biosynthesis and cell cycle modules, whereas the VIM lineage was enriched in the EMT module. Conversely, POU2F3 and NEPC cell types exhibited enrichment in the NE2 module, which displayed lineage-specific biological functions across different subpopulations (Figure 2F). The signature of EMT module was associated with a worse prognosis based on RNA-seq data from TCGA (Figure S2C). Consistent with the enrichment of inflammatory signaling pathways in the VIM lineage, we observed significant positive correlations between the EMT module and NF-κB signaling (Figure 2G). These observations characterized a unique VIM lineage exhibiting a hybrid of mesenchymal and stem-like traits with activated NF-κB signaling.

**Figure 2:**
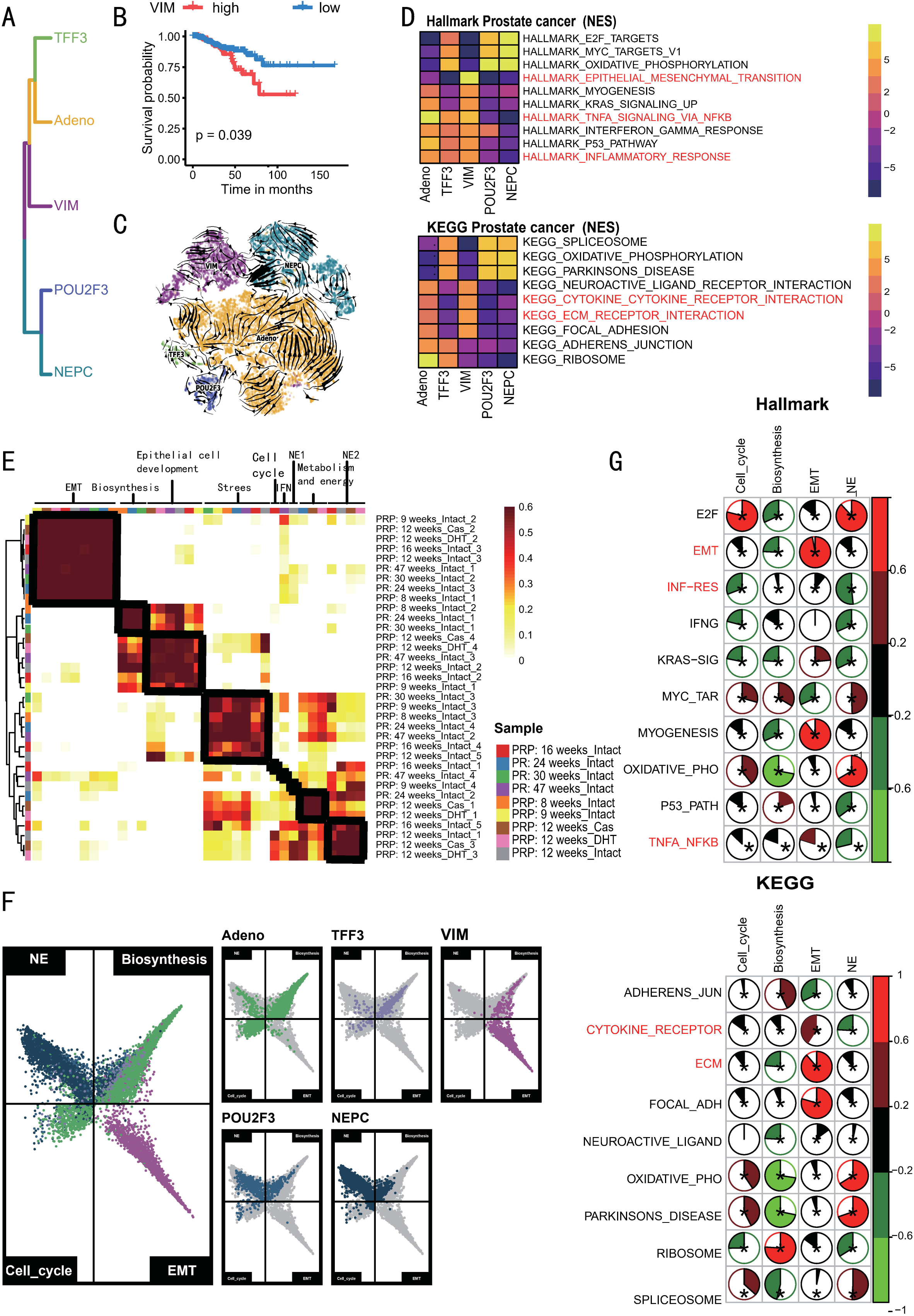
Molecular heterogeneity of five malignant lineages. (**A**) Unsupervised hierarchical clustering for five malignant lineages. (**B**) Survival probability of patients with prostate cancer (from TCGA data) stratified by expression levels of the VIM lineage gene signature. (**C**) RNA velocity vectors of five malignant lineages on t-distributed stochastic neighbour embedding. (**D**) Heatmap showing the highest two enriched pathways in malignant lineages based on Hallmark Pathway gene sets (above) and KEGG Pathway gene sets (below). (**E**) Pairwise correlation clustering of 35 gene expression programs (GEPs) of malignant cells with forming nine consensus modules whose biological function predicted by GO analysis (top). (F) Two-dimensional butterfly plot visualization of the NE, biosynthesis, Cell_cycle and EMT modules in five malignant lineages, representing modules scores as relative meta-module scores. Each quadrant corresponds to one modules; the exact position of each cell reflects its relative signature scores in all four modules. (**G**) Correlogram showing Pearson correlation between top differentially enriched pathways (top: Hallmark Pathway; below: KEGG Pathway) and four modules (NE, biosynthesis, Cell_cycle and EMT modules). Asterisks indicating statistically significant comparisons (*P*-value[<[0.05) and scale bars indicating Pearson correlation (r) (red[=[positive correlation, green[=[negative correlation).

### Infiltration of GRN^+^ macrophage is associated with lineage transition

During the lineage transition, Mono/Macro/DC composition underwent significant changes (Figure 1D). To explore the interactions between TME compositions and malignant cells, we then inferred the receptor-ligand (R-L) interactions using CellPhoneDB[30]. Our analysis revealed that Mono/Macro/DC were the main cell types that secrete ligands targeting malignant cells, especially VIM lineage (**Figure 3A**), indicating that Mono/Macro/DC might play a crucial role in shaping the VIM lineage. To investigate the relationship between different cell types in prostate cancer, we utilized CIBERSORTx to predict the abundance of cell subtypes in TCGA cohort (https://portal.gdc.cancer.gov/), DKFZ-PRAD cohort (https://www.cbioportal.org/), and other public bulk RNA-seq and microarray datasets[31–34]. Spearman correlations were performed to examine the infiltration patterns of the 10 predominant cell types across six independent prostate cancer cohorts. We found a notable positive correlation between Mono/Macro/DC subsets and VIM lineage in the majority of cohorts (Figure 3B). Furthermore, we found significant positive correlations between of Mono/Macro/DC signature and plasticity features, specifically the VIM lineage signature (Figure S2D). Notably, the abundance of Mono/Macro/DC cells showed a significant increase in CRPC patients compared with adenocarcinoma patients and controls (Figure S2E). To assess the clinical significance of immune cell infiltration within TME, we analyzed the association between Mono/Macro/DC infiltration and disease-free survival (DFS) in patients from TCGA and Dkfz cohorts. Mono/Macro/DC infiltration was significantly associated with worse DFS in these patient cohorts (Figure S2F). These findings indicate that Mono/Macro/DC are key regulators of malignant cell states and contributors to VIM lineage development within the TME.

**Figure 3:**
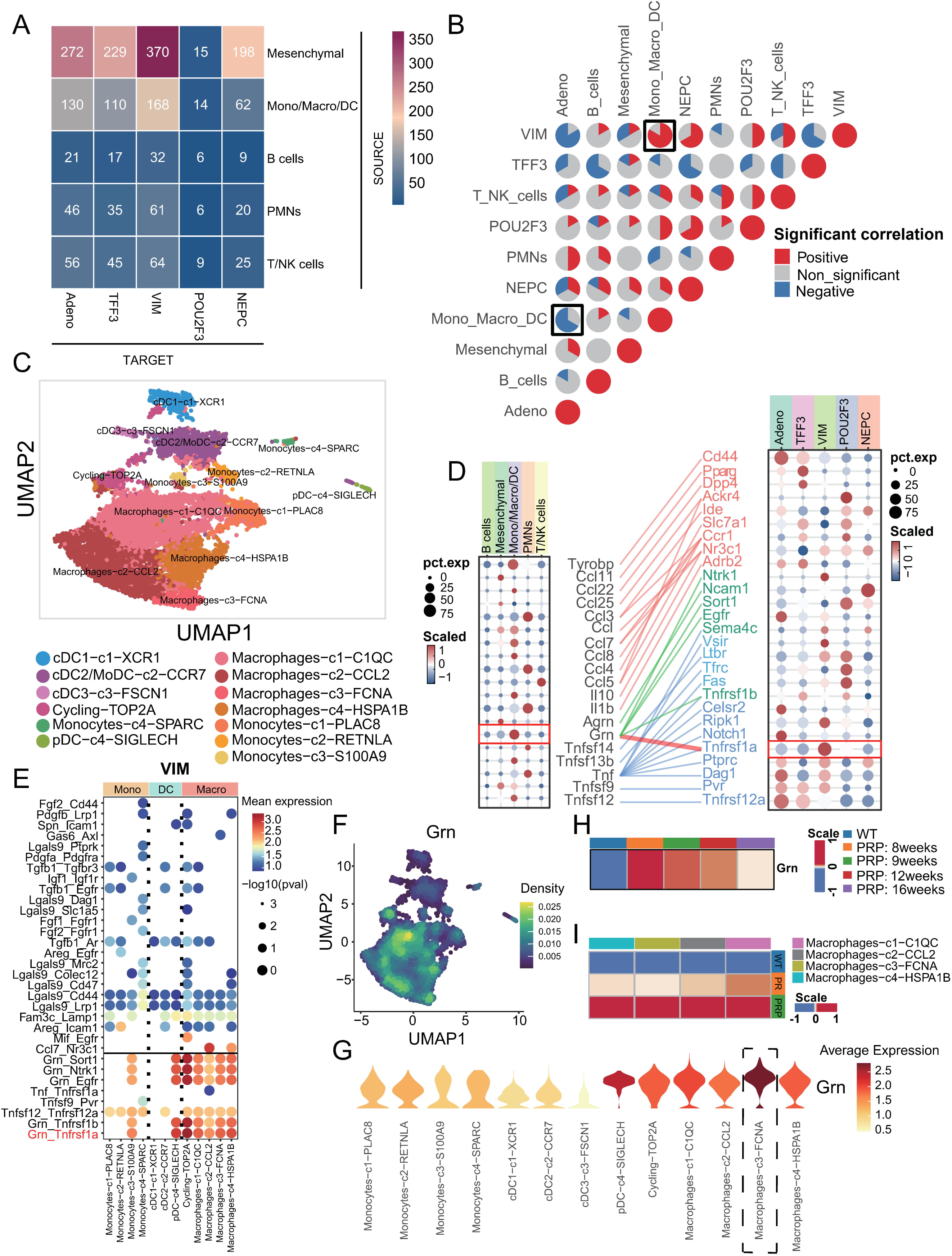
GRN^+^ macrophage promotes lineage transition for prostate cancer. (A) Heatmap showing the number of receptor-ligand interactions between tumor microenvironment cell types and five malignant lineages. (B) The proportion of PRAD cohorts exhibiting significant positive (Spearman correlation with Rs > 0.2 and FDR < 0.05, shown in red), negative (Rs < -0.2 and FDR < 0.05, shown in blue), or non-significant (gray) correlations in the infiltration levels between pairwise cell types across six independent PRAD cohorts. (C) UMAP visualization depicts cell populations colored by annotated Mono/Macro/DC subsets. (D) Dot plots showing selected ligand-receptor pairs for interactions of five malignant lineage and Mono/Macro/DC subsets. Dot size indicates the *P* value generated by permutation test, and color indicates the mean expression of each ligand-receptor pair. (E) Dot plots showing selected ligand-receptor pairs for interactions of VIM lineage and Mono/Macro/DC subsets. Dot size indicates the *P* value generated by permutation test, and color indicates the mean expression of each ligand-receptor pair. (F) UMAP visualization depicts expression density of Grn in Mono/Macro/DC. (G) Violin plots showing the expression level of Grn across Mono/Macro/DC subsets. (H) Heatmap showing Grn gene expression scores in Mono/Macro/DC at different timepoints. (I) Heatmap showing Grn gene expression scores in Macrophage subsets at different sample types.

We re-clustered and categorized Mono/Macro/DC into 13 sub-populations (Figure 3C, Figure S3A-C). Compared with WT mice, the composition of Mono/Macro/DC in PtRP mice exhibited a positive correlation with the early progression stage, with expansion occurring in PRP_8weeks_Intact (Figure S3D). We then performed DEG analysis of the paracrine inflammation genes mentioned earlier[35] between Mono/Macro/DC in PRP:8-9weeks_Intact and WT, as well as between PRP:12-16weeks_Intact and PRP:8-9weeks_Intact. Interestingly, we found that some paracrine genes, such as Spp1 and Grn were exclusively expressed by Mono/Macro/DC during the lineage transition stage and declined upon termination of reprogramming (Figure S4A). High expression of *SPP1* is associated with vascular-related genes and has been reported to be significantly correlated with poor survival prognosis in multiple types of tumors[36–38]. Granulin, encoded by *Grn* gene, is secreted by inflammatory cell in responses to wounds and is involved in tumorigenesis and tumor progression[39]. Nielsen et al.[40] and Quaranta et al.[41] demonstrated that granulin fosters the activation of the mesenchymal program, resulting in immune checkpoint resistance and metastasis in pancreatic cancer. Consistent with the results of the DEG analysis, the expression of *Grn* in Mono/Macro/DC was upregulated during lineage progression and downregulated at terminal stages (Figure 3H), suggesting that granulin may play a role in driving EMT programs and the development of VIM lineage. Furthermore, we assessed the strengths of ligand-receptor interactions between Mono/Macro/DC subsets and five malignant lineages using CellPhoneDB[30] (Figure 3E, Figure S4B). Some ligand-receptor pairs showed strong signaling from Mono/Macro/DC subsets to the VIM lineage, including Grn_Sort1, Grn_Ntrk1, Grn_Egfr, Tnfsf9_Pvr, Tnfsf12_Tnfrsf12a, Grn_Tnfrsf1b, and Grn_Tnfrsf1a(Figure 3E). We focused on Grn, Tnfsf12, and Tnfsf9 as potential ligands. To identify key ligands, we first scored EMT, stemness, and NF-κB activity in malignant cells and averaged these scores per sample. We then measured and averaged ligand expression in Mono/Macro/DC cells for each sample, and finally assessed the correlations between ligand levels and malignant features. *Grn* showed strong correlations with stemness (0.52), EMT (0.42), and NF-κB signaling (0.44), while *Tnfsf12* and *Tnfsf9* had weak correlations (R < 0.3) (Figure S4C). This observation identified Grn as the primary ligand driving the malignant state among the candidates. Consistent with the CellPhoneDB result, Mono/Macro/DC expressed the highest levels of GRN expression among TME compositions (Figure 3D, Figure S4D). Meanwhile, the Tnfrsf1a receptor was predominantly expressed in the VIM lineage (Figure 3D, Figure S4E). Furthermore, to identify the myeloid subpopulation that exhibits high expression of *Grn*, we observed a high expression of *Grn* in macrophages (Figure 3F). Notably, *Grn* was primarily expressed in FCNA^+^ macrophages (Figure 3G), which showed MDSC-like characteristics and played a tumor-promoting role in colon cancer[38], liver cancer[42] and lung cancer[43]. In addition, *Grn* was predominantly expressed at high levels in macrophages of the PRP group (Figure 3I), where the tissues in the group exhibited a higher degree of malignancy compared with PR and WT group, suggesting macrophages secrete granulin to fuel cancer progression in highly malignant microenvironments. These data emphasize the important role of macrophage-derived granulin in driving the malignant lineage state.

We next analyzed the trajectories of CRPC-Adeno and NEPC to delineate dynamic signaling scores along the lineage transition state. Scores of plasticity features, such as stemness and EMT, and the expression of Tnfrsf1a was the highest in the intermediate stage (Figure S5A). In particular, TNFA_SIGNALING_VIA_NFKB signaling exhibited the highest expression score in the intermediate stage (Figure S5A). Furthermore, we computed inflammatory signaling and lineage transition signaling scores for each malignant cell and segmented them into 39 clusters. The TNFA_SIGNALING_VIA_NFKB signaling showed a strong positive correlation with plasticity feature scores and *CD44* expression score, but a significant negative correlation with AR signature score (Figure S5B). NF-κB signaling has been reported to activate genes to control cancer stemness and motility change[44], and correlates with poor survival for patients with prostate cancer[45, 46]. Other pathways including TGF-β signaling pathway (25), Wnt/β-catenin signaling pathway (26), Notch signaling pathway (27), and Hippo/YAP signaling pathway (28), have been reported to promote EMT. To assess their potential contributions, we performed pathway scoring on the VIM lineage across GEMMs, human prostate cancer (described below), and TRAMP (described below) samples. Our results showed that the NF-κB signaling pathway had the highest score, while the scores for the other pathways were relatively lower, reinforcing the dominant role of NF-κB signaling in the VIM lineage in our study (Figure S5C). These results suggest that macrophage-derived granulin promotes VIM lineage formation by activating NF-κB signaling

### GRN^+^ macrophage-derived granulin mediates EMT program and drives AR-targeted therapy resistance

To verify whether granulin can enhance the cellular viability of RM-1 cells, RM-1 cells were treated with secreted recombinant mouse Grn protein for 24h, 48h and 72h, respectively. CCK8 assay showed that the OD scores were significantly higher in RM-1 cells treated with recombinant mouse Grn protein compared to control cells (*P* < 0.01, **Figure 4A**). We then conducted co-culture assay of RAW264.7 cells and RM-1 cells in a non-contact Transwell system that produce a cell culture environment that resembles an in vivo state and allow cells to carry out metabolic activities in a more natural manner without cell-to-cell contact (Figure 4B). Then, to examine whether Grn^+^ macrophage could promote EMT program of RM-1 cells, Grn overexpression plasmid was constructed and transfected into mouse macrophage RAW264.7 cells, RT-qPCR confirmed Grn mRNA was significantly overexpressed in RAW264.7 cells compared with control cells (Figure 4C,Left). Remarkably, RM-1 co-cultured with Grn^+^ macrophage became stretched and elongated compared with control RM-1 cells (Figure 4F). As shown in Western blot, Grn^+^ macrophage dramatically suppressed epithelial marker E-cadherin and ZO-1, and upregulated mesenchymal marker Vimentin (Figure 4D). Conversely, lentiviruses carrying shRNAs against Grn were transfected into RAW264.7 cells, RT-qPCR confirmed lentivirus-mediated Grn shRNA significant downregulated Grn mRNA in RAW264.7 cells (Figure 4C, Right), consequently Grn^-^ macrophage increased epithelial protein E-cadherin and ZO-1, but depressed mesenchymal marker Vimentin and N-Cad (Figure 4D). Taken together, these macrophage-derived granulin facilitated EMT program in prostate adenocarcinoma cells. To explore the signaling pathways activated in RM-1 cells when co-cultured with Grn^+^ macrophages, bulk RNA sequencing was performed. Consistent with previous findings in VIM lineage (Figure 2D, Figure S5), GSEA analysis indicated a notable activation of inflammatory pathways in RM-1 cells co-cultured with GRN^+^ macrophages, with top enrichment of the NF-κB signaling pathway (Figure 4E), In summary, GRN^+^ macrophage may elicit EMT and morphological transformation via activating NF-κB signaling in prostate adenocarcinoma cells.

**Figure 4:**
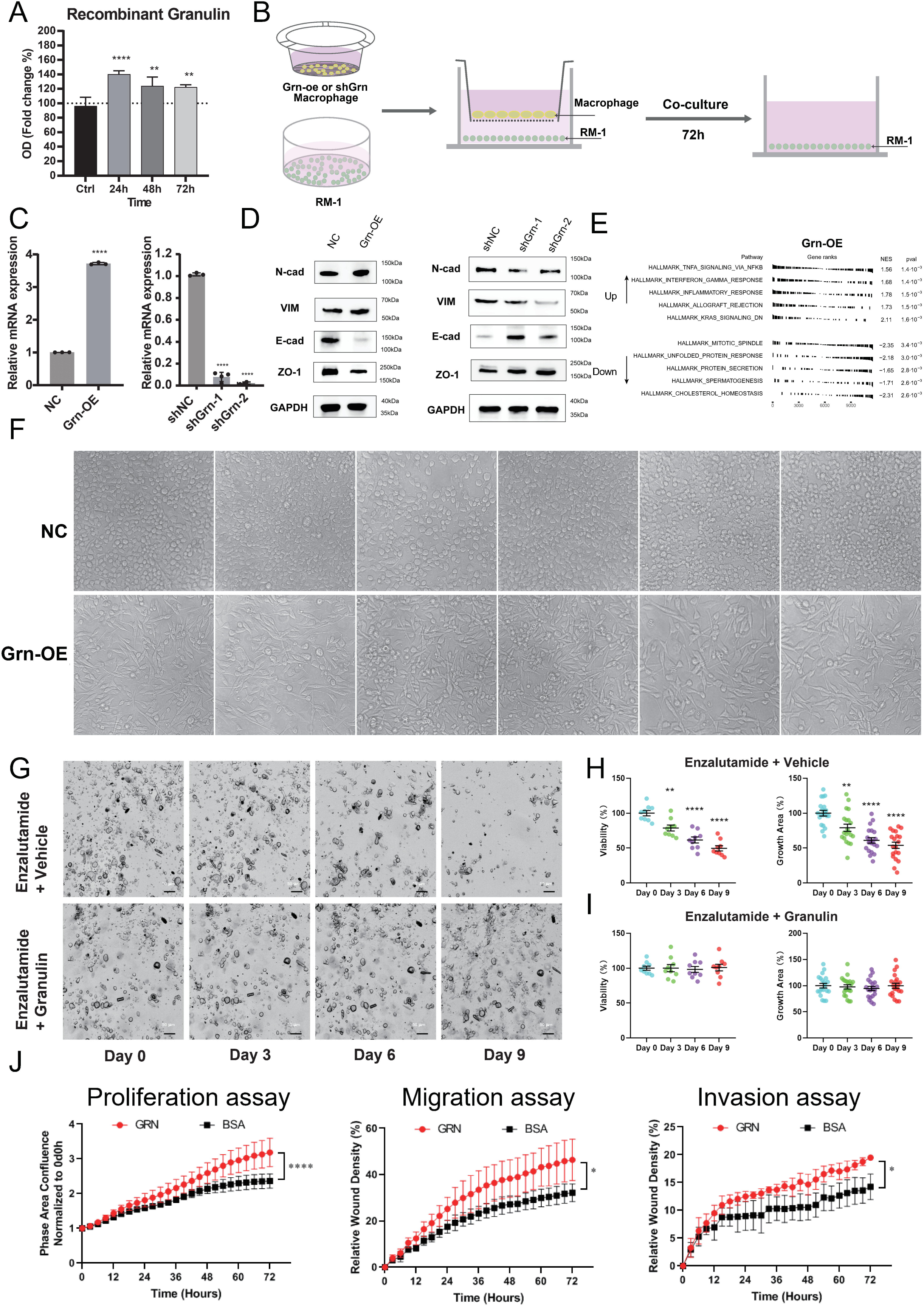
Granulin augment EMT and proliferation of prostate cancer cells in vitro. (**A**) Cell viability of RM-1 cells treated with indicated doses of recombinant granulin. Data measured using CCK8 assays. Data are expressed as the mean±SD, \*\**P*<0.01, \*\*\*\**P*<0.0001 by one-way ANOVA. (**B**) Schematic illustration of co-culture macrophage RAW264.7 cells with RM-1 cells. (**C**) Quantitative RT-PCR of overexpressing the GRN gene (left) or shRNA targeting Grn gene (right) in mouse macrophage RAW264.7. Bars represent fold change in mRNA levels relative to NC. Data are expressed as the mean±SD, \*\**P*<0.01, \*\*\*\**P*<0.0001 by one-way ANOVA. **(D)** Immunoblots of indicated EMT proteins in RM-1 cells post co-culture with RAW264.7 transfected with overexpressing the Grn gene (left) or shRNAs (right). (E) fGSEA analysis showing the up-regulated and down-regulated enriched pathways for RM-1 cells post co-culture with RAW264.7 with overexpressing the Grn. ranked by P-value. **(F)** Morphology of RM-1 cells post co-culture with RAW264.7 with overexpressing the Grn (n=6). (**G**) Representative images of patient-derived prostate cancer organoids after enzalutamide (80 μM) treatment with or without granulin (16.6 μg/mL). Scale bars: 50 μm. **(H, I)** Patient-derived prostate cancer organoid viability (n = 9 wells from 3 patients) and growth area (n = 20 fields from 3 patients) after enzalutamide (80 μM) treatment with (below) or without (above) granulin (16.6 μg/mL). Viability was measured using CCK8 assay. Growth area was measured using high-content assay. Date are mean±s.d, \*\**P*<0.01, \*\*\*\**P*<0.0001 by one-way ANOVA. (**J**) Live-cell imaging assay of LNCaP cells with or without recombinant human granulin protein to measure cell growth, migration and invasion over 3 days. Date are mean±s.d, \*\**P*<0.01, \*\*\*\**P*<0.0001 by two-way ANOVA.

To determine if the granulin can promote AR-targeted drugs resistance in prostate cancer patients, we established three patient-derived prostate cancer organoids. We utilized recombinant granulin protein and a clinical-grade AR signaling inhibitor (enzalutamide) to investigate the functional consequences of granulin on AR-targeted drug efficacy. Our results demonstrated that enzalutamide substantially inhibited the growth and activity of organoids (Figure 4G-I). In contrast, granulin promote resistance to enzalutamide, maintaining both their quantity and activity (Figure 4G-I). This revealed the potential role of granulin in enhancing resistance to AR-targeted drugs.

To determine if the granulin can promote the metastasis. We used the human prostate cancer cell line LNCap to perform proliferation, migration and invasion assays, and found that granulin significantly promotes LNCap cell proliferation, migration, and invasion (Figure 4J, Figure S6). This showed potential role of granulin in contributing to metastasis and tumor progression.

### Crucial TME compositions and lineage programs are confirmed in human CRPC

To investigate the relationship between transcriptional changes and the progression of plasticity in human prostate cancer, we re-analyzed scRNA-seq data from 3 CRPC-Adeno patients and 4 CRPC-NEPC patients in a previous study[23]. We divided single cells that passed all quality control filters into twenty clusters and manually annotated into five known cell types (**Figure 5A**, Figure S7A). As expected, myeloid cells in the human samples showed specific expression of *GRN* (Figure 5C). Malignant cells were then extracted and clustered into 13 distinct clusters (Figure S7B), which were further annotated into three major cell types: malignant epithelium, mixed lineage, and NEPC (Figure 5B). The mixed lineage cells exhibited specific expression of receptor TNFRSF1A (Figure 5C), and high expression of several EMT, stemness and NE genes, including *VIM, TGFB2, KLF4, CD44, FOXA2, GHGB* and *POU3F2* (Figure 5C, Figure S7B), indicating their location in a mixed lineage state with multilineage potential. Meanwhile, the mixed lineage signature was associated with unfavorable patient survival based on RNA-seq data from TCGA (Figure S9A). Consistent with our findings from scRNA-seq of GEMMs, we observed a significant positive correlation between the abundance of myeloid cells and mixed lineage in most patient cohorts (Figure S8A).

**Figure 5:**
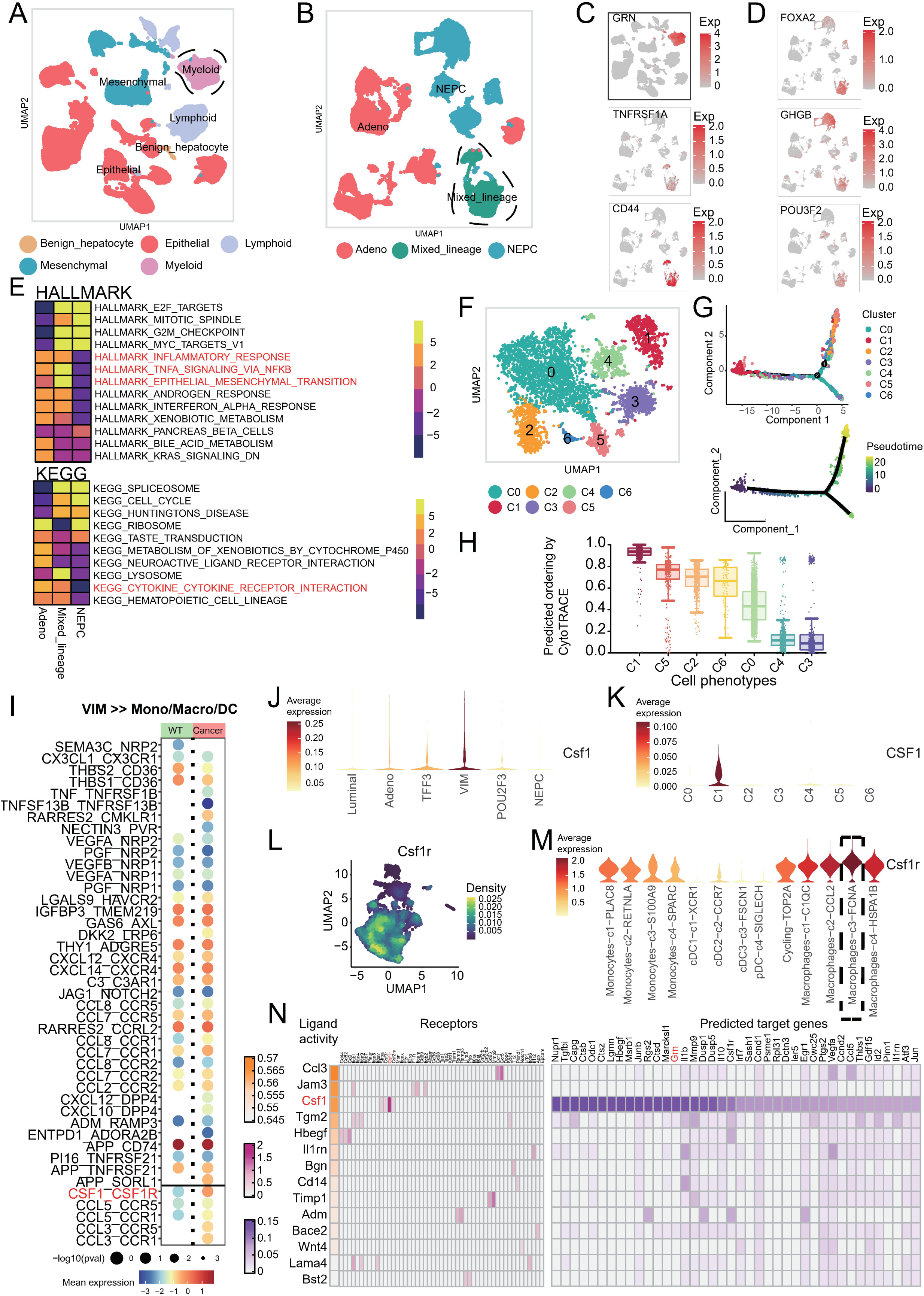
Crucial TME compositions and lineage programs arises in CRPC patient biopsies. (A) UMAP visualization depicts all cells from CRPC patient biopsies colored by annotated major cell types. (B) UMAP visualization depicts tumor cells from CRPC patient biopsies colored by annotated tumor cell types. (C, D) UMAP visualization of GRN, TNFRSF1A, CD44 gene markers (C) and FOXA2, GHGB and POU3F2 (D). (E) Heatmap showing the top enriched pathways in malignant lineages based on GSEA Hallmark Pathway gene sets (above) and KEGG Pathway gene sets (below). (F) UMAP visualization depicts cell populations of mixed lineage from CRPC patients biopsies colored by clusters. (G) Developmental trajectory of mixed lineage colored by mixed lineage clusters (above) and pseudotime scores (below). (H) Boxplot showing the CytoTRACE score of mixed lineage clusters. (I) Dot plots showing selected ligand-receptor pairs for interactions from VIM lineage to Mono/Macro/DC among WT and GEMMs mice. Dot size indicates the *P* value generated by permutation test, and color indicates the mean expression of each ligand-receptor pair. (J) Violin plots showing Csf1 expression level across Luminal cells and five malignant lineage cells from GEMMs. (K) Violin plots showing CSF1 expression level across mixed lineage clusters from CRPC patient biopsies. (L) UMAP visualization depicts expression density of Csf1r in Mono/Macro/DC from GEMMs biopsies. (M) Violin plots showing the expression level of Csf1r across Mono/Macro/DC subsets from GEMMs biopsies. (N) Heatmap showing top-ranked ligands inferred to regulate Mono/Macro/DC by VIM lineage from GEMMs biopsies using NicheNet.

Myeloid cells were also positively correlated with plasticity features and specifically the VIM lineage signature (Figure S8B). Additionally, the abundance of myeloid cells was significantly increased in CRPC patients (Figure S8C), which correlated with poorer overall survival in patients from both the TCGA and Dkfz cohorts (Figure S8D). Besides, we found that inflammatory responses and EMT hallmarks were significantly enriched in mixed lineage (Figure 5E). Employing Monocle3, we inferred lineage states and confirmed that mixed lineage was an intermediate lineage based on pseudotime ordering (Figure S7B, C). The proportion of mixed lineage cells initially increased but subsequently declined, while the fraction of NEPC cells steadily rose, and the proportion of Adeno cells gradually decreased along the lineage progression (Figure 7D). Meanwhile, we found that TNFRSF1A expression and the scores of AR, NEPC, EMT, stemness, and TNFA/NF-κB signatures exhibited a consistent trend with our findings in GEMMs, along with pseudotime ordering (Figure S7E). The highest levels of EMT, stemness, and TNFA/NF-κB program scores were observed in the mixed lineage, while the scores of AR and NEPC signatures in the mixed lineage exhibited intermediate levels between malignant epithelium and NEPC (Figure S7C). Furthermore, we computed TNFA/NF-κB signaling, EMT, and stemness signature for malignant cells and segmented them into 56 clusters. Scatterplots of these clusters showed a strong positive correlation between TNFA/ NF-κB signaling and EMT, as well as between TNFA/ NF-κB signaling and stem-like traits. (Figure S7D). These findings were consistent with our observations from scRNA-seq of GEMMs, suggesting that EMT program and NF-κB signaling play a crucial role in lineage transition in human CRPC.

### Granulin-NFκB-CSF1 axis contributes to macrophages-induced lineage transition

To investigate the origin of human mixed lineage, we performed further clustering of the mixed lineage and identified seven distinct cell clusters (Figure 5F). Using monocle2[47] and CytoTRACE[48], we confirmed that C1 cluster exhibited the highest stemness and represents the origin of the mixed lineage (Figure 5G, H). We scored the human mixed lineages and observed that C1 cluster exhibited hybrid mesenchymal and stem-like characteristics with highly activated NF-κB signaling, which was similar to VIM lineage from GEMMs, suggesting this cell state possesses the potential for multilineage redifferentiation into other lineages (Figure S10A). Interestingly, C1 cluster was enriched in cytokine production and chemotaxis by GO analysis (Figure S10B), indicating this activated NF-κB signaling in this cell state might induce the production of cytokines and chemokines that attract and educate inflammatory cells to sustain tumor-promoting TME[49].

We performed CellphoneDB analysis to determine what cytokines in the VIM lineage was released to educate the macrophages in the TME. And we found that VIM lineage interacted more frequently with macrophages in tumor group compared with WT group, where CSF1_CSF1R and several chemokine receptor interactions, such as CCL3_CCR5 and CCL3_CCR1 were significantly enriched in tumor group (Figure 5I). CCL3 has been reported to recruit macrophage[50, 51] while CSF1 can promote M2 macrophage polarization[52]. Consistently, VIM lineage showed high expression level of *Csf1* among malignant and luminal cells in GEMMs (Figure 5J). Meanwhile, the expression level of *CSF1* was also highly expressed in C1 cluster among mixed lineage in human CRPC (Figure 5K). We next investigated the receptor *Csf1r* expression in Mono/Macro/DC composition and found that *Csf1r* was mainly expressed in macrophages (Figure 5L), especially in the FCNA^+^ macrophages (Figure 5M) that highly expressed *Grn* (Figure 3G), suggesting FCNA^+^ macrophages with high *GRN* expression were the main receptor cells which may respond to over-expressed ligand CSF1 from malignant cells.

We further investigated the key regulatory molecules and downstream activated targets from the VIM lineage to macrophages. Using NicheNet[53], we found that the VIM lineage exhibited high Csf1 ligand activity to bind to Csf1r receptors, resulting in the elevated expression of target gene *Grn* in macrophages (Figure 5N). Moreover, the top 100 predicted targets in macrophages were enriched in TNF signaling pathway, IL-17 signaling pathway and NF-κB signaling pathway (Figure S10C). IL-17 signaling and NF-κB signaling have previously been reported to induce M2-like phenotype[54–56]. Consistently, treatment of RAW264.7 cells with the Csf1 ligand resulted in morphological changes indicative of altered polarization. RT-qPCR analysis confirmed that Csf1 induced M2 polarization, which was accompanied by an upregulation of GRN expression(Figure S10D). Taken together, these findings indicated that there is a positive feedback loop between GRN^+^_FCNA^+^ macrophages and CSF1^+^ VIM lineage, which promotes the malignant lineage transition and the recruitment and polarization of macrophages.

### Pharmacologic blockade of the CSF-1/CSF-1R axis suppresses VIM lineage formation

Based on our findings, we next wonder whether disrupting the positive feedback loop can inhibit prostate cancer lineage transition. We administered a small molecule inhibitor of CSF1R (BLZ945) to treat TRAMP mice for 7 days. Then, vehicle-treated and BLZ945-treated prostate tumor tissue were collected for scRNA-seq (**Figure 6A**). We classified the transcriptomes into 10 major cell types based on canonical gene markers (Figure 6B; Figure S11A). Using ClusterFoldSimilarity[57] and Qiming Zhang et al.’s method[42], we computed similarity scores of cell clusters and we found the VIM lineage from GEMMs and TRAMP mice showed extremely high similarities (Figure 6C, Figure S11B). To confirm whether the VIM lineage originated from malignant cells, Adeno, and VIM lineage were re-clustered into 13 clusters whose large-scale chromosomal copy number variation (CNV) was calculated using Infercnv[58]. We found that VIM lineage and all the Adeno clusters exhibited significantly higher CNV levels than random control cells (Figure S11D), suggesting the VIM lineage may be transitioned from malignant cells in these TRAMP mice. Of note, the signature of the VIM lineage was significantly associated with poor clinical prognosis based on RNA-seq data from TCGA (P=0.03, Figure S11E). Compared with Adeno, VIM lineage showed remarkable up-regulation of genes associated with myeloid growth factors (*Csf1*), chemokines (*Ccl2*) and mesenchymal markers (*Vim, Col1a2, Col3a1*) (Figure S11F). The biomarker genes of VIM lineage was enriched in GO terms like inflammatory response, angiogenesis and interferon response (Figure 6D). Moreover, VIM lineage harbored mesenchymal and stem-like traits with activated NF-κB signaling (Figure S11G). Taken together, the VIM lineage in the TRAMP mice shares the similar biological features as same as these in the GEMMs.

**Figure 6:**
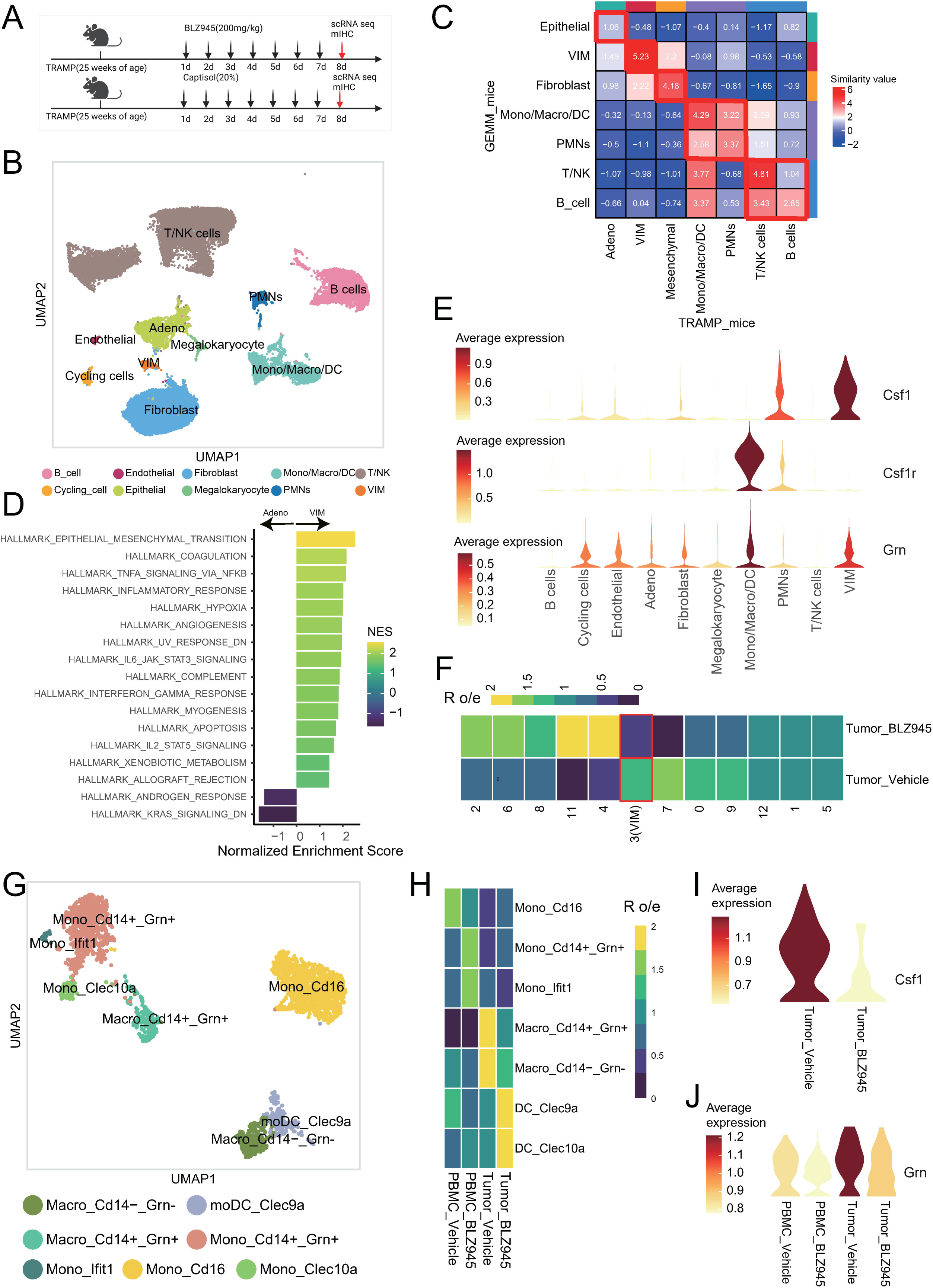
Pharmacologic blockade of CSF-1 suppresses VIM lineage formation in TRAMP mice models. (**A**) Experimental design schematic of BLZ945 on TRAMP mice at the 25 weeks of age. Following 7-day treatment, prostate tumors were collected for scRNA-seq and mIHC, respectively. (**B**) UMAP visualization depicts cell populations colored by annotated major cell types of TRAMP mice. (**C**) Heatmap showing the similarities of cell clusters of GEMMs mice (x-axis) and TRAMP mice(x-axis) using ClusterFoldSimilarity. (**D**) Gene set enrichment analysis (GSEA) (Hallmark Pathway gene sets) on genes ranked by log2 fold change between VIM lineage and Adeno. NES, normalized enrichment score. (**E**) Violin plots showing the expression level of Csf1(above), Csf1r (middle) and Grn (below) across major cell types in TRAMP mice. (**F**) Heatmap showing tissue preference of cell subsets of Adeno clusters and VIM lineage across prostate tumor tissues from TRAMP mice treated with BLZ945 and vehicle by Ro/e. (**G**) UMAP visualization depicts Mono/Macro/DC cell populations colored by annotated major cell types of TRAMP mice. (**H**) Heatmap showing tissue preference of Mono/Macro/DC cell populations across prostate tumor tissues and blood samples from TRAMP mice treated with BLZ945 and vehicle by Ro/e. (**I**) Violin plots showing the expression level of Csf1 in VIM lineage across prostate tumor tissues from TRAMP mice treated with BLZ945 and vehicle. **(J)** Violin plots showing the expression level of Grn in Mono/Macro/DC cell populations across prostate tumor tissues and blood samples from TRAMP mice treated with BLZ945 and vehicle.

As expected, Csf1 was mainly expressed in the VIM lineage among all cell clusters (Figure 6E, top). TRAMP mice treated with BLZ945 showed a reduction of VIM lineage enrichment with a decrease of *Csf1* expression (Figure 6F, I). As anticipated, Mono/Macro/DC expressed *Csf1r* and *Grn* at the highest level among all cell clusters (Figure 6E, middle and bottom). Mono/Macro/DC were re-clustered into 7 clusters (Figure 6G), and BLZ945 reduced the Grn+ macrophages enrichment in tumor tissues (Figure 6H) with decrease of *Grn* expression (Figure 6J). These results indicated that BLZ945 may disrupt the positive regulatory loops, leading to a decreased *Grn* expression of macrophages, thereby suppressing VIM lineage formation.

### Mass Cytometry analysis of tissues from patients with CRPC-Adeno and CRPC-NEPC

To validate our findings at single-cell protein level, we enrolled CRPC patients (n=3) and NEPC patients (n=4) and collected tumor tissues for Cytometry by the time of flight (CyTOF) (**Figure 7A**). Low-quality cells were excluded using Cytobank (www.cytobank.org) and the remaining cells were clustered using FlowSOM based on 41 parameter panels. CyTOF analysis identified 10 major cell types visualized by t-SNE projection (Figure 7B, Figure S12). A mixed lineage was identified that highly expressed EMT and stemness markers, including *Snail.1*, *Vimentin*, *ZEB2* and *OCT3-4* (Figure 7C), which was similar to the VIM lineage and mixed lineage from scRNA-seq of GEMMs (Figure 3) and human CRPC (Figure 5), respectively. In addition, cell proportion of Mono/Macro and mixed_lineage showed an increase in NEPC patients (Figure 7D). Moreover, the expression of adenocarcinoma epithelium (EpCAM) and NE (Synatophys) marker in the mixed lineage exhibited intermediate levels compared with that in malignant epithelium and NEPC, but the expression of EMT markers (Snail.1, Vimentin, ZEB2) and stemness markers (OCT3-4) were highest in the mixed lineage (Figure 7E). Notably, NF-κB activation factor NFkB.p65 was significantly highly expressed in mixed_lineage (Figure 7F), and granulin was highly expressed in Mono/Macro (Figure 7G). In brief, these observations were consistent with the results from scRNA-seq data analysis.

**Figure 7:**
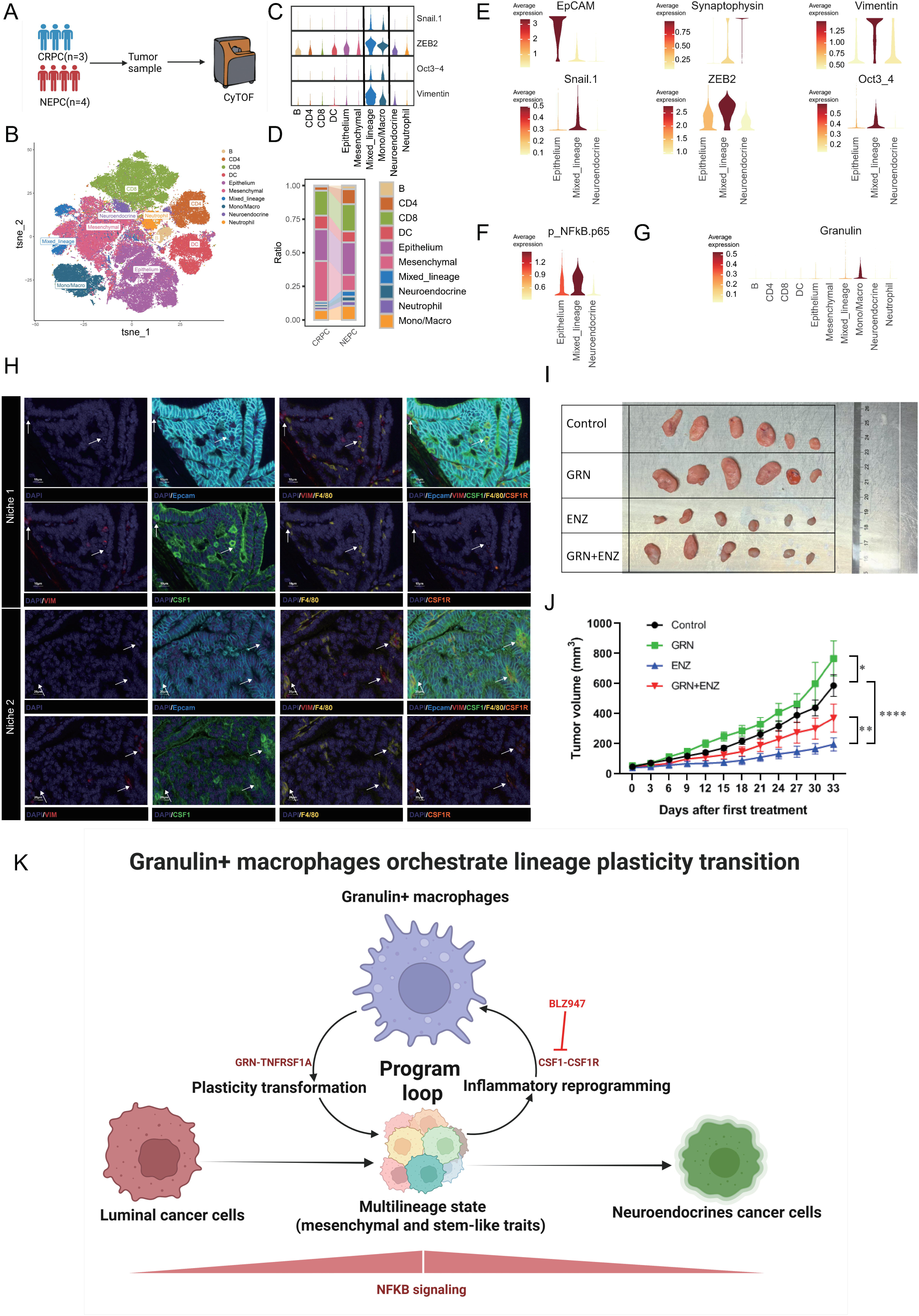
Mass cytometry analysis of phenotypic and compositional heterogeneity of cells from CRPC patients biopsies. (**A**) Schematic representation of CyTOF experimental design. (**B**) UMAP projection of cells from CRPC-Adeno (n=3) and CRPC-NEPC (n=4) patient biopsies colored by FlowSOM clusters. (**C**) Violin plot showing signal intensity of Snail.1, ZEB2, Oct3-4, Vimentin across ten major clusters. (**D**) Fraction of cell types (y-axis) of CRPC-Adeno and CRPC-NEPC patient biopsies colored by annotated ten cell types. (**E**) Violin plot showing signal intensity of elected markers across epithelium, mixed lineage and neuroendocrine cells. (**F**) Violin plot showing signal intensity of NFkB.p65 across epithelium, mixed lineage and neuroendocrine cells. (**G**) Violin plot showing signal intensity of granulin across ten major clusters. (**H**) Multiplex immunofluorescence staining to show the co-localization of VIM^+^ cells and macrophage (F4/80^+^). (**I**) Tumor tissues were dissected 33 days after inoculation. (**J**) Time-dependent growth of tumor tissues in xenograft model. Date are mean±SEM, n=6, \*\**P*<0.05, \*\**P*<0.01, \*\*\*\**P*<0.0001 by two-way ANOVA. (**K**) Proposed model system of granulin+ macrophages orchestrating the multilineage state with mesenchymal and stem-like traits. Panel created BioRender. Huafeng, F. (2025) https://BioRender.com/w4rie22.

To further validate our findings at the spatial level, we obtained spatial transcriptomics data from tissue have been defined as NEPC coexisting with HSPC[59]. We classified these cells into 5 clusters based on gene markers, AR signature and NEPC signature (Figure S13A, C). Marker genes of tumor-associated macrophages, including MSR1, CSF1R, and CD163, showed high expression in the NEPC_TME area. (Figure S13B). Additionally, *GRN* expression, scores of GRN^+^ Mono/Macro, TNFA_SIGNALING_VIA_NFKB and stemness signatures were enriched in NEPC_TME, exhibiting significant spatial co-localization (Figure S13B, D). We further performed multiplex immunofluorescence staining of tumor samples from TRAMP mice and validated that macrophages were spatially close to VIM+ cells in many niches (Figure 7H). In contrast, we observed that such interactions were almost nonexistent in benign tissue (Figure S13E). In summary, these data suggested GRN^+^ macrophages and VIM lineage were enriched in the NEPC microenvironment, contributing to the transformation-induced plasticity.

We then conducted in vivo experiments using xenograft model, granulin (16.6 μg/mL) treatment significantly promoted tumor growth compared to BSA group (p < 0.05). Treatment with enzalutamide (ENZ; 10 mg/kg, every other day) effectively suppressed tumor progression (p < 0.0001). However, granulin (16.6 μg/mL) co-treatment significantly impaired the efficacy of ENZ, resulting in significantly larger tumors compared to ENZ treatment alone group (p < 0.01)(Figure 7I, J), suggesting that granulin could contribute to resistance to AR-targeted therapy.

## Discussion

It is well-known that the cell state in tumor phenotype is influenced by a combination of cell-intrinsic factors (genetic and epigenetic alterations)[13–19, 60–62] and cell-extrinsic (autocrine, paracrine) factors[12, 23, 63]. In this study, we revealed notable alterations in the cellular composition in TME and multiple malignant lineages transition during the progression of prostate cancer. We discovered a pivotal population of GRN-positive macrophages that secrete granulin, leading to the transformation of tumor cells into a multilineage state by activating NF-κB signaling. Subsequently, the multilineage state induces macrophages to express elevated levels of GRN through the release of CSF1, establishing a positive feedback loop in cellular communication (Figure 7K).

Currently, the majority of reported drivers implicated in prostate cancer lineage plasticity are not suitable for pharmacological targeting[64]. Here, we identified granulin as a ligand that drives lineage transition and AR-targeted therapy resistance. Granulin is a pluripotent growth factor that regulates tumor progression[65], embryonic development[66] and wound healing[39]. In terms of malignancy, granulin has been reported to regulate stem cell and cancer stem cell signaling molecules[67, 68], such as Oct4, Nanog, Sox2, and CD133, resulting in the acquisition of stem cell traits and a poorer prognosis for HCC patients[69]. Granulin has been reported in multiple studies that it can promote tumor progression and metastasis, including in liver cancer[70], breast cancer[71], and pancreatic cancer[40]. In addition, macrophage-secreted granulin can initiate EMT program to promote cancer metastasis[40] and drive resistance to immune checkpoint therapy[41]. However, its potential role in facilitating lineage plasticity remains largely uncertain. Our results revealed that granulin modulates NF-κB signaling activity in tumor cells. Consequently, NF-κB signaling activation plays a role in transitioning to a stem-like and EMT state[44] but not in directly redifferentiating to NE-like lineage. Furthermore, the special malignant lineage upregulates CSF1 expression that is required for M2 macrophage polarization[52] and promotes granulin production[72] in a positive feedback fashion. The CSF-1/CSF-1R axis blockade in TRAMP mice validates the crucial role of the positive feedback loop’s integrity in the process of lineage transition.

Multiple cell types work synergistically to promote lineage transition and tumor progression. In addition to Mono/Macro/DC subsets, our analysis revealed that the mesenchymal component played a crucial role in the TME (Related Manuscript File). We identified a specialized subset of endothelial cells, DCN^+^ endothelial, which possess characteristics of EndMT. These cells with EndMT trait play a coordinating role in malignant tumor proliferation and promote metastasis[73]. Regarding fibroblasts, a crucial stromal component known as CCL7^+^ fibroblast, which highly express SPRP1, displayed a strong distribution preference within tumor tissue. It has been reported that SFRP1 is involved in driving an epithelial NE differentiation transition-like phenotype, contributing to therapy resistance and metastatic progression in prostate cancer[74]. The functions of polymorphonuclear subtypes in tumorigenesis been extensively studied[63, 75–77], while the involvement of neutrophils in prostate cancer is still mostly unknown. We identified a pro-tumor neutrophil subpopulaiton, IFIT1^+^ neutrophil, which spectrum highly expresses PDL1 and exhibits an immunosuppressive phenotype. Similarly, Zhang *et al.* reported a similar pro-tumor neutrophil in live tumor studies[78]. This finding provides a promising target for immunotherapy in prostate cancer.

The TRAMP and GEMM models were applied in our study, which share core molecular features and conserved regulatory pathways with human prostate tumors[79]. The two models effectively recapitulate the progression of human prostate cancer, including the critical adenocarcinoma-to-neuroendocrine transformation observed in clinical patients[80, 81], and have been widely used in many studies to model disease progression and deterioration[23, 82].

However, there are some limitations. Firstly, the TRAMP and GEMM mice represent different experimental mouse models. However, our cell similarity analysis demonstrates that cell clusters annotated as the same cell subsets show high similarities between the two mouse models. Additionally, the VIM lineage exhibits similar activated pathways and highly expressed genes between the two mouse models. Further inferCNV analysis revealed VIM lineage in TRAMP mouse model exhibited high copy number variation scores, suggesting such VIM lineage undergoes lineage transition from the adenocarcinoma akin to GEMMs in TRAMP mice.

In summary, our study provides insight into a previously undiscovered mechanism involving the formation of a multilineage state via the reprogramming of a special kind of macrophages. It also elucidates the critical roles of the granulin-NF-κB signaling-CSF1 loop in mediating lineage transition, potentially offering promising therapeutic targets aimed at overcoming AR-targeted therapy resistance.

## Materials and methods

### Collection and acquisition of mouse models and human prostate tumors

Prostate tumor samples utilized for Cytometry by Time-Of-Flight (CyTOF) analysis were obtained from 3 CRPC-Adeno and 4 CRPC-NEPC patients who underwent transurethral prostatectomy at Shanghai Tongji Hospital. Written informed consent was provided by each patient. The clinical characteristics of the patients included in this study are shown in Table S1. TRAMP mice (male, 25 weeks, weighing 40 g) for pharmacologic blockade were purchased from Jiangsu Huachuang Xinnuo Pharmaceutical Technology Co., Ltd.

Single-cell RNA expression profiles of 29 mice samples (9 wildtype [WT], 7 Pten-/-Rb-/- [PtR], and 13 Pten-/-Rb-/-Trp53-/- [PtRP]) and 12 human prostate cancer samples were collected from GSE210358 by Joseph M. Chan[23]. Spatial transcriptome sequencing data was obtained from GSE230282[59]. Additionally, RNA-seq data and clinicopathological data from various sources were utilized, including The Cancer Genome Atlas (TCGA) PRAD cohort (483 prostate patients), GSE54460 (106 prostate patients)[32], GSE116918 (248 prostate patients)[83], GSE54460 (107 prostate patients)[32], GSE70768 (13 CRPC tissues, 113 tumour tissues, 73 matched benign tissues)[34], GSE70769 (93 prostate patients)[34], GSE94767 (154 prostate cancer patients)[33] and DKFZ-PRAD (268 prostate patients). These datasets were obtained from the TCGA database (https://portal.gdc.cancer.gov/), cBioPortal (https://www.cbioportal.org/) and GEO Datasets (https://www.ncbi.nlm.nih.gov/gds/).

### scRNA-seq analysis for the genetically engineered mouse models

A total of 103,684 single cells from the genetically engineered mouse models (GEMMs) were initially integrated. To ensure data quality, potential doublet cells were identified and removed using Scrublet[84]. Three quality filteration was conducted for the integrated dataset: 1) The total UMI count per cell was required to be above 500. 2) The number of detected genes per cell was required to be above 250 and below 6000. 3) The percentage of mitochondrial genes was required to be below 20%. Based on these criteria, 83,183 cells were retained for downstream analysis using R package Seurat v4[85]. After selecting 1,500 highly variable genes, principal component analysis was performed to calculate 50 principal components (PCs). Nearest neighborhood graphs were constructed using the top 50 PCs for clustering based on the FindNeighbors function. Then, a total of 15 major cell types were annotated using well-known marker genes by the FindClusters function. The marker genes used for annotation included *HOXB13*, *SBP*, *KRT4*, *FOXI1*, *SDC1*, *CALML3*, and *PATE4* for normal epithelial cell lineage (L1, L2, L3, Basal, Basal SV, SV); *KRT14*, *KRT5*, *EPCAM*, *TFF3*, *VIM*, *POU2F3*, *CHGA*, and *GFP* for malignant cell lineage (Adeno, TFF3, POU2F3, VIM, NEPC); *CD79A*, *MS4A1*, *CD3E*, and *CD3D* for lymphoid cells (B, NK_T); *COL1A1* and *COL1A2* for mesenchymal cells (endothelial cells and fibroblasts); *CD68* and *CST3* for myeloid cells (Mono_Macro_DC), *CD14* and *CSF3R* for myeloid cells (PMNs). To further annotate the cell subtypes, a second round of clustering was performed on the major cell types respectively.

### scRNA-seq analysis for human prostate cancer patients

The pre-processing of single-cell sequencing data for human prostate cancer samples followed the same methodology as previously described in single-cell RNA-seq analysis for GEMM samples. Five major cell types were annotated based on well-known marker genes, including *CLDN5*, *COL1A1*, and *VIM* for mesenchymal cells; *CD14*, *LYZ*, and *PTPRC* for myeloid cells; *CD2*, *CD3E*, and *CD3D* for lymphoid cells; *AR*, *CDKN2A*, *EPCAM*, and *KLK3* for epithelial cells; *ALB* and *CRP* for benign hepatocytes. Malignant cells were identified and further re-clustered into three sub-cell types using the same procedure as the initial round. These subtypes include *AR* and *KLK3* for malignant epithelial cells, *NOTCH1*, *KLF4*, *TGFB2*, *VIM*, and *SMAD2* for mixed lineage cells, and *ASCL1*, *CHGA*, and *SYP* for NEPC cells.

### scRNA-seq analysis for TRAMP mouse models

TRAMP mouse models can progress to poorly differentiated prostate cancer with NE characteristics by 24 weeks[86]. For TRAMP mice at 25 weeks of age, CSF-1R inhibitor BLZ945 was orally administered daily at a concentration of 200 mg/kg and captisol (20%) was served as the vehicle control for 7 days. Fresh prostate tissues were collected for single-cell transcriptome sequencing. Library construction and sequencing were performed at Shanghai OE Biotech. Co. Ltd. The pre-processing of sequencing data for TRAMP mice samples followed the same methodology as previously described in single-cell RNA-seq analysis for GEMM samples. Ten major cell types were annotated based on well-known marker genes, including *Cd79a* and *Mzb1* for B cells; *Cd3d*, *Cd3e*, and *Nkg7* for T/NK cells; *Epcam* for Adeno; *Cd14* and *Cd68* for Mono/Macro/DC; *S100a8* and *S100a9* for PMNs; *Gp9* and *Pf4* for Megalokaryocyte; *Top2a* for Cycling cells; *Col1a1* and *Col1a2* for Fibroblast; *Pecam1* for Endothelial; *Vim* for VIM. Adeno and VIM were identified and further re-clustered into 13 sub-cell types, Mono/Macro/DC was further re-clustered into 7 sub-cell types. The procedure was the same as the first round. To assess cell type enrichment across tissues, we compared observed and expected cell numbers within each cluster using the previously defined formula[87].

### Pathway enrichment analyses

We employed three different approaches to evaluate enriched pathways. 1) Gene ontology (GO) analysis: uniquely differentially expressed genes specific to relevant groups, clusters, or cell types were identified using the ‘‘FindAllMarkers’’ function R package Seurat as described above. These genes underwent GO enrichment analysis using gene sets obtained from the Gene Ontology database (http://www.geneontology.org/). The clusterProfiler package version 21 was utilized for this analysis[88]. 2) Gene set enrichment analysis (GSEA): For the enrichment analysis of relevant groups, clusters, or cell types, we utilized Hallmark gene sets (RRID: SCR_016863) and Kyoto Encyclopedia of Genes and Genomes (KEGG) gene sets (https://www.genome.jp/kegg/). The GSEA[89] (http://software.broadinstitute.org/gsea/index.jsp) was employed to identify pathways with FDR < 0.05 as significantly enriched. 3) Gene set signature scoring: For single-cell transcriptome data, several published signatures were obtained to score cells using AUCell[23, 90–95]. For spatial transcriptomic data, GRN^+^ mono/macro is the designation for cells with gene expression levels of *GRN* higher than the mean among mono/macro in GEMMs, and its signature genes were obtained using COSG[86]. The gene signature score for each cell was computed by mean expression across all genes included in the signatures.

### Survival analysis for novel subclusters in bulk RNA-seq data

To analyze the survival outcomes of novel subclusters within bulk RNA-seq data, the following steps were carried out: signature specific to different cell populations, such as Fibroblast-c1-CCL7, Endothelial-c2-ACKR1, Endothelial-c3-DCN, PMN-c6-IFIT1, VIM lineage in GEMM, mixed lineage in human prostate cancer patients, were selected based on specific criteria, including a fold-change greater than 2, adjusted *P*-values below 0.01, and a cell expression percentage higher than 0.1. A signature score of each cell population was determined as the mean expression of signature genes based on RNA sequencing data from TCGA, and then divide the patients into two groups based on the median of signature score. The two-sided long-rank test was used to compare Kaplan-Meier survival curves.

### Cell infiltration analysis based on single-cell data

To estimate infiltration of our defined 10 major cell types in GEMM samples and human CRPC patients samples from TCGA cohort (https://portal.gdc.cancer.gov/), DKFZ-PRAD cohort (https://www.cbioportal.org/) and other public bulk RNA-seq and microarray datasets[31–34], we generated a reference signature matrix of major cell types using CIBERSORTx[96]. We used a permutation parameter setting of 500 repetitions to impute cell fractions while keeping all other parameters at their default values. Spearman correlation analysis was performed among cell-type infiltration levels to assess cell infiltration correlation, and a correlation coefficient greater than 0.2 with a false discovery rate (FDR) below 0.05 was considered statistically significant. To explore the prognostic value of cell population infiltration, we divided them into two groups based on the median infiltration of each cell type from the TCGA cohort (https://portal.gdc.cancer.gov/) and the DKFZ-PRAD cohort (https://www.cbioportal.org/). The two-sided long-rank test was used to compare Kaplan–Meier survival curves.

### Assessment of the association between NF-κB signaling and lineage plasticity

For single-cell transcriptome data from GEMMs, we calculated the inflammatory signaling and lineage transition signature scores for each malignant cell, then segmented them into 39 clusters with a resolution of 2 using the “FindClusters” function in Seurat. Spearman’s correlation analysis was performed to assess the relationship between inflammatory signaling and lineage transition signature scores for these clusters. For single-cell transcriptome data from human CRPC patients samples, we calculated the TNFA_SIGNALING_VIA_NFKB signaling, EMT and stemness signature scores for each malignant cell, then segmented them into 56 clusters based on a resolution of 1.2 using the “FindClusters” function in Seurat. Spearman’s correlation analysis was performed to assess the relationship between TNFA_SIGNALING_VIA_NFKB signaling and EMT and stemness.

### Assessment of the association between myeloid signature and lineage plasticity

Mono/Macro/DC signature from GEMM and myeloid signature from human CRPC patients were selected based on specific criteria, including a fold-change greater than 2, adjusted *P*-values below 0.01, and a cell expression percentage higher than 0.1. We computed the mean expression of signature genes to obtain the signature score based on RNAseq data from TCGA. Spearman correlation was performed between Mono/Macro/DC or myeloid signature scores and lineage plasticity signature scores.

### Developmental trajectory analysis

The developmental trajectory was inferred using Monocle2 (v.2.12)[47]. Differentially Expressed Genes (DEGs) were identified using the differential Gene Test function, and genes with a q-value of less than 1×10^-5^ were employed to order the cells based on pseudotime. Subsequently, we employed the DDRTree algorithm to perform dimension reduction and arrange the cells along the trajectory. Monocle3 was performed to map the cell fate[97]. The expression data underwent UMAP embedding using the Monocle function ’reduce_dimension’ with default parameters. Subsequently, the trajectory graph was inferred using the “learn_graph” function, setting the minimal branch length to 15 and the “close_loop” to FALSE. In addition, The Slingshot software was utilized to infer the lineage trajectory[98].

### cNMF analysis

It is well-known that a single cell contains specific cell function programs. We applied the cNMF algorithm to obtain consensus modules representing various cell programs. The algorithm principles were based on a previous study[99]. To obtain the best solution, we performed the cNMF algorithm 100 times and selected the optimal value of k, which ranged from 5 to 15, for each sample. As a result, we identified 9 gene expression programs (GEPs) in PRP_16weeks_Intact, 4 in PR_24weeks_Intact, 3 in PR_30weeks_Intact, 4 in PR_47weeks_Intact, 3 in PRP_8weeks_Intact, 3 in PRP_9weeks_Intact, 4 in PRP_12weeks_Intact, 4 in PRP_12weeks_DHT, and 5 in PRP_12weeks_Intact. We then calculated a gene-set score for each GEP in all malignant cells within each sample. Next, we performed correlation clustering of the cross-sample GEP scores. The ‘1 - Pearson correlation coefficient’ was used as the distance metric and “ward D2” linkage, which allowed us to identify consensus modules. To represent each GEP, we selected the top 50 genes based on their differential expression. GO analysis of the top 50 genes of each GEP was conducted to further characterize these modules. The mean expression of the top 50 genes for the EMT program based on RNA-seq data from TCGA was divided into high and low risk groups. The two-sided long-rank test was used to compare Kaplan-Meier survival curves.

### Cell communication analysis

Cell communication analysis was performed to study cell-cell interactions across Mono/Macro/DC subsets and malignant cells in GEMM samples using CellPhoneDB package[100]. The significance of cell-cell communication was determined based on the interaction and the normalized cell matrix obtained via scran normalization, where a significance threshold of *P* < 0.05 was considered as statistically meaningful. To pinpoint potential ligands secreted by VIM lineage contributing to the macrophage in tumor samples from GEMM samples, we performed NicheNet[53] to evaluate the regulatory network of VIM lineage on macrophage. Macrophages in the samples from WT were considered as reference receiver cells.

### RNA velocity analysis

Utilizing the Python package velocyto[101], we computed RNA velocity values for each gene in every cell, mapped the RNA velocity vectors into a low-dimensional space, visualized these vectors on a UMAP projection.

### Copy number variation (CNV) analysis

The initial CNV value of each individual Adeno and VIM cell was estimated using infercnv[99], where randomly selected endothelial cells and fibroblasts were used as reference cells. The initial CNV values were transformed based on the following criteria, followed by calculating the average CNV for each chromosome:

- For values where [CNV values>0.85 & CNV values<0.9], set to 2.
- For values where [CNV values>=0.9 & CNV values<0.95], set to 1.
- For values where [CNV values>=0.95 & CNV values<1.05], set to 0.
- For values where [CNV values>=1.05 & CNV values<1.1], set to 1.
- For values where [CNV values>=1.1 & CNV values<1.15], set to 2.

### Mass Cytometry

Tumor samples from 7 CRPC patients (3 CRPC-Adeno, 4 CRPC-NEPC) were collected for Mass Cytometry analysis and the clinical information was showed in Supplementary Table 1. Live/dead staining was performed using 2 μM cisplatin (Fluidigm) followed by quenching with CSB (Fluidigm). Subsequently, cells were first stained and fixed with Fix-I buffer (Fluidigm), then secondarily stained using the Cell-ID™ 20-Plex Pd Barcoding Kit (Fluidigm) to reduce sample cross-reactivity. Antibodies were labeled with Maxpar® x8 Polymer Kits (Fluidigm) as the manufacturer’s protocol. For surface marker labeling, cells were counted, diluted to 1×10^6 cells/ml, and and incubated with a surface antibody cocktail. Following this, cells were washed, permeabilized with 80% methanol, and stained with an intracellular antibody cocktail. Following triple washes with CSB, cells were incubated with 0.125 μM iridium intercalator in fix and perm buffer (Fluidigm) overnight at 4 [. Before acquisition, samples were washed and resuspended in deionized water containing 10% EQ 4 Element Beads (Fluidigm), with cell concentrations adjusted to 1×10^6 cells/ml. Data were acquired on a Helios mass cytometer (Fluidigm). The raw FCS files were normalized and compiled per sample.. Subsequently, all .fcs files were uploaded to Cytobank for data cleaning, and populations of single living cells were exported as .fcs files for further analysis.

For CyTOF data processing and analysis, we performed initial manual gating to identify live, intact, single cells and then applied boolean gating for de-barcoding using Cytobank (www.cytobank.org). Cells meeting the criteria were downloaded as FCS files and compensated for arcsinh transformation (scale factor 5) for subsequent analysis. Major clusters were generated with FlowSOM/Consensus ClusterPlus within CATALYST[102, 103] and visualized using UMAPs and the ComplexHeatmap package[104].

### Spatial transcriptomic data processing

The spatial transcriptome data of tissue diagnosed as NEPC coexisting with HSPC[59] was normalized and adjusted using the Sctransform method, following the tutorial manual provided by Seurat (https://satijalab.org/seurat/articles/spatial_vignette.html). GRN^+^ Mono/Macro cells in GEMM were defined as macrophages expressing GRN gene at levels greater than the mean expression value. The top 20 marker genes of GRN^+^ Mono/Macro were defined as GRN^+^ Mono/Macro signature using COSG[2]. Signature scores were computed by averaging the expression levels of all signature genes. Spearman correlation between GRN and TAM markers (CD163, MSR1, CSF1R) was calculated using the function “cor()” in R.

### Cell and organoid culture

Mouse prostate cancer RM-1 cells and Human prostate cancer LNCaP cells were obtained from Procell Life Science & Technology Co., Ltd (Wuhan, China) and maintained with RPMI 1640 supplemented with 10% fetal bovine serum (FBS) plus antibiotics (Invitrogen, USA). RAW264.7 and HEK293T cells were obtained from National Infrastructure of Cell Line Resource (Beijing, China) and cultured in Dulbecco’s Modified Eagle Medium (DMEM) media supplemented with 10% FBS in an incubator at 37°C with 5% CO_2_, 100 % humidity. Prostate tumor samples utilized for organoid culture were obtained from 3 patients who underwent radical prostatectomy at Shanghai Tongji Hospital. Written informed consent was provided by each patient. Human prostate cancer organoids were generated and cultured according to the manufacturer’s instructions and maintained in human prostatic carcinoma organoid media (Cat#PRS-OCR-100, Cat#PRS-ODR-100, Cat#PRS-TDD-2-100, Cat#PRS-TCR-1-100, Cat#PRS-TDE-2-100, Cat#PRS-PRCM-3D-100, Precedo. Matrigel Cat#356231, BD).

### Cell and organoid viability

CCK8 assay was used to measure the cell viability. Briefly, cells seeded in 96-well plates were treated with ligand recombinant protein (recombinant mouse TNFSF12, Cat#50174-M15H, Sino Biological; recombinant mouse IL-1β Protein, Cat#50101-MNAE, Sino Biological; secreted recombinant Mouse GRN Protein, Cat#50396-M08H, Sino Biological) for the indicated concentrations and durations, respectively. Organoids seeded in 24-well plates were treated with enzalutamide (Cat#T6002, TargetMol) with or without recombinant Human Granulin Protein (Cat#10826-H08H, Sino Biological). Cell and organoid viability was determined using a CCK-8 kit (Cat#B34302, Selleck) according to the manufacturer’s instructions.

### High-content imaging and analysis

Assay data and images were acquired on an Operetta CLS high-content imaging and analysis system (PerkinElmer, Inc., Waltham, MA) with a 5x air objective. Six fields were acquired per well. All images were analyzed using Harmony 5.2 software (PerkinElmer) to calculate growth area.

### shRNA knockdown and over-expression of ligand genes

Hairpin shRNAs targeting the coding sequences (CDS) of mouse Tnfsf12, Il-1b, and Grn transcripts were inserted into the pLL3.7 vector (Plasmid Cat#11795, Addgene). The specific targeting sequences used for each hairpin are provided in Supplementary Table 2. Over-expression sequences of the ligands are listed in Supplementary Table 3. To package lentiviral particles, HEK293T cells were co-transfected with 10 μg of the pLL3.7 shRNA construct or the overexpress construct, 2 μg of VSVG, and 5 μg of PAX2. The transfection was performed using Lipofectamine 3000 (Cat#L3000015, Thermo Fisher) according to the manufacturer’s instructions. At 72 hours post-transfection, the supernatant containing the viruses was collected and filtered through a Filter Unit with a 0.45 μm pore size and a PVDF membrane (Millipore). The viruses were then concentrated by centrifugation at 18000 rpm, 4°C for 2 hours. The resulting sediment was resuspended using DMEM culture media.

### RNA extraction and real-time PCR analysis

Total RNA was isolated using an RNA extraction kit (Cat#DP451, TIANGEN) and reverse transcribed using TransScript® All-in-One First-Strand cDNA Synthesis SuperMix (Cat#AT341-01, TransGen) according to the manufacturer’s protocol. Quantitative PCR was performed in triplicates with SYBR Green PCR master mix (Cat#A57156, Thermo fisher scientific), and the sequence of primers used are listed in Supplementary Table 4.

### Co-culture of macrophage with RM-1 cells

The treated mouse macrophage RAW264.7 cells were seeded in transwell inserts with 0.4[μm pore size (Cat#3412, Corning) with the RM-1 cells cultured in 6-well plates at a ratio of 1:1. The transwell inserts containing RAW264.7 cells were removed after 72[h of co-culture and the RM-1 cells were collected for subsequent analysis.

### Immunoblots

Cells were harvested and homogenized in stringent-RIPA lysis buffer containing a protease inhibitor cocktail (Cat#B14001, Selleck) for 20 min on ice and followed by centrifugation at 12000 rpm, 4[for 10 min. The collected cell supernatants were suspended in an SDS loading buffer and incubated at 100°C for 10 min. The primary antibodies used were rabbit anti-TNFRSF1A (Cat#21574-1-AP, Proteintech,1:1000), rabbit anti-N-cadherin (Cat#13116, Cell signaling technology, 1:1000), rabbit anti-Vimentin (Cat#5741, Cell signaling technology,1:1000), rabbit anti-E-cadherin (Cat#3195, Cell signaling technology, 1:1000), rabbit anti-ZO-1 (Cat#8193, Cell signaling technology, 1:1000), rabbit anti-GAPDH (Cat#5174, Cell signaling technology,1:5000). Secondary antibodies conjugated to HRP were used at 1: 5000.

### Proliferation, migration and invasion assay

LNCaP cells were seeded at a density of 4–5 × 10³ cells per well in 96-well plates. Real-time monitoring of cell proliferation was performed using the IncuCyte S3 live-cell imaging system (Sartorius, Essen BioScience, USA), with images automatically captured every 3 hours over a 3-day period without removing the cells from the incubator. Cell confluence was quantified using the IncuCyte Base Software (v2024A). Each experiment was conducted in five replicates per condition. For migration and invasion assays, cells were plated at a density of 4 × 10[cells per well in 96-well plates. A uniform scratch wound was created using the 96-well WoundMaker tool once the monolayer reached confluence. For invasion assays, Matrigel matrix (Corning, #356231) was added to each well. Time-lapse images and quantitative data for migration and invasion were automatically acquired every 3 hours over a period of 3 days using the IncuCyte Base Software (v2024A).

### Animals and treatments

All animal experiments were performed according to the animal protocols approved by the Institutional Committee of Peking Third Hospital. Male immunodeficient BALB/c nude mice, aged 6–8 w, were purchased from Beijing Vital River Laboratory Animal Technology Co., Ltd, China. Approximately ∼10^7^ LNCaP cells pretreat with GRN (16.6 μg/mL) or BSA in a 200 μl suspension of a 1:1 mixture of medium and Matrigel (Corning, 356,231) were subcutaneously injected into the flanks of mice. The length and width of the tumor were measured every 3 days, and the tumor volume (mm^3^) was estimated using the following formula: (longest diameter) × (shortest diameter)^2^ × 0.5. Enzalutamide (ENZ, 10 mg/kg) was given in oral administration every other day (n = 6 per group).

### Multiplex immunofluorescence staining

Multiplex immunofluorescence staining for Epcam, CSF1, F4/80, Granulin and Vimentin was conducted on prostate cancer tissues from TRAMP mice using the Opal 7-Color Automation IHC Kit (Cat#NEL821001KT; Akoya Biosciences). Briefly, 5-μm thick formalin-fixed, paraffin-embedded (FFPE) sections were stained in sequence using a reference protocol[105]. All the antibodies and their respective fluorophore were as follows: Epcam (1:500, Dye Opal 480, Cat#ab213500, Abcam), F4/80 (1:200, Dye Opal 570, Cat#70076, Cell Signaling Technology), CSF1 (1:300, Dye Opal 520, Cat#ab233387, Abcam), CSF1R (1:100, Dye Opal 620, Cat#ab313648, Abcam) and Vimentin (1:200, Dye Opal 690, Cat#5741, Cell Signaling Technology). The multispectral images from stained slides were scanned by Motif model on the PhenoImager HT system (Akoya Biosciences) at high magnification (40x, 0.25 μm/pixel).

## Ethical statement

This study was approved by the Ethics Committee of Tongji Hospital affiliated to Tongji University (KYSB-2023-105). This study was conducted in accordance with the principles of the Declaration of Helsinki, and written informed consent was obtained from all patients prior to enrollment.

## Data availability

Raw sequencing datasets of RNA-seq and scRNA-seq reported in this paper have been deposited at the Genome Sequence Archive (GSA) in the National Genomics Data Center, China National Center for Bioinformation, Chinese Academy of Sciences with accession number CRA019064 and are publicly accessible at https://ngdc.cncb.ac.cn/gsa/browse/CRA019064.

## CRediT authorship contribution statement

**Zhipeng Zhu:** Methodology, Investigation, Validation, Software, Formal analysis, Writing-original draft, Writing – review & editing, Data curation. **Mi Zhang:** Methodology, Investigation, Validation. **Ying Song:** Methodology, Investigation, Validation, Funding acquisition. **Wei Jiang:** Methodology, Investigation. **Fang Cao:** Data curation, Validation. **Yicong Yao:** Data curation, Resources. **Xiaotong Yu:** Validation. **Hongyu Zhao**: Data curation. **Husile Baiyin**: Data curation. **De Chang**: Resources. **Xiaolu Zhao**: Supervision. **Gang Wu:** Conceptualization, Supervision, Resources, Writing – review & editing. **Kailong Li:** Conceptualization, Supervision, Resources, Writing – review & editing, Funding acquisition. **Fengbiao Mao:** Conceptualization, Supervision, Resources, Funding acquisition, Writing – review & editing.

## Supporting information

Related Manuscript File

## Acknowledgements

We appreciate the professional suggestions from Prof. Shudong Zhang in Department of Urology, Peking University Third Hospital.

