## Supplemental Tables for "Granulin^+^ macrophages promote lineage plasticity in prostate cancer through paracrine signaling loops"

**Supplementary Table 1**

Clinical information of tumor samples from 7 CRPC patients (3 CRPC-Adeno, 4 CRPC-NEPC) were collected for Mass Cytometry analysis

| **Experimental sample identifier** | **Sample type** | **Age** | **Gleason score** | **Body weight（Kg）** | **PSA（ng/mL）** |
| --- | --- | --- | --- | --- | --- |
| A8_MFB01_H01_T012 | CRPC-Adeno | 76 | 7 | 66 | 2000 |
| A8_MFB01_H01_T002 | CRPC-Adeno | 66 | 8 | 66 | 100 |
| A8_MFB01_H01_T018 | CRPC-Adeno | 87 | 10 | 60 | 101 |
| A8_MFB01_H01_T011 | CRPC-NEPC | 80 | 9 | 65.8 | 4.2 |
| A8_MFB01_H01_T001 | CRPC-NEPC | 76 | 8 | 85 | 3.2 |
| A8_MFB01_H01_T017 | CRPC-NEPC | 73 | 6 | 50 | 9.15 |
| A8_MFB01_H01_T016 | CRPC-NEPC | 78 | 9 | 60 | 5.2 |

**Supplementary Table 2**

Sequences for shRNAs targeting ligands in macrophage

| shRNA | Targeting sequence |
| --- | --- |
| shGRN-1 | CCTAGAATAACGAGCCATCAT |
| shGRN-2 | ACTCATCCTGAGTCACCCTAT |

**Supplementary Table 3**

Sequences for overexpression of ligands in macrophage

| Gene | Forward sequence | Reverse sequence |
| --- | --- | --- |
| Grn | TGGGTCCTGATGAGCTGGCTGG | CAGTAGCGGTCTTGGGACCGGA |

**Supplementary Table 4**

Primers sequence for RT-qPCR

| Gene | Forward | Reverse |
| --- | --- | --- |
| Grn-1 | CTGTTGCCTGCTGCCTTGAC | ACACCCTTAGAGAACGGGCAG |
| Grn-2 | TGTAAGGAAGGGCTACAGAC | CCACAGAAACCGGAAGAAATG |
| Il-4 | ATCATCGGCATTTTGAACGAGG | TGCAGCTCCATGAGAACACTA |
| Il10 | GCTCTTACTGACTGGCATGAG | CGCAGCTCTAGGAGCATGTG |
| Tgfβ | CCACCTGCAAGACCATCGAC | CTGGCGAGCCTTAGTTTGGAC |
| Gapdh | AGGTCGGTGTGAACGGATTTG | TGTAGACCATGTAGTTGAGGTCA |
