## Supplementary material for "Granulin^+^ macrophages promote lineage plasticity in prostate cancer through paracrine signaling loops": Related Manuscript File

**A special fibroblast subset is associated with** **malignant lineage transition**

Mesenchymal cells were isolated and re-clustered into four subtypes: endothelial, fibroblast, lymphatic endothelia, and oligodendrocyte-like stromal cells (Supplementary Figure 14A). We re-clustered fibroblast cells into 10 subtypes (Supplementary Figure 14B, C, D, F). One specific subset, fibroblast-c1-CCL7, showed significant alterations in PR and PRP (Supplementary Figure 14E), consistent with a strong distribution preference observed in PtR and PtRP based on odds ratio analysis (Supplementary Figure 14H). Importantly, the gene signature of fibroblast-c1-CCL7 was significantly associated with a worse prognosis (Supplementary Figure 14G), highlighting its pivotal role in immune microenvironment deterioration. In PtRP, we identified thousands of DEGs specific to fibroblast-c1-CCL7 during different stages of cancer progression, with significant upregulation observed at PtRP_8weeks_Intact (Supplementary Figure 14I). Interestingly, DEGs analysis revealed the enrichment of SFRP1 in PtRP_8weeks_Intact (Supplementary Figure 14J). The composition of fibroblast-c1-CCL7 and the SFRP1 expression exhibited an increasing trend at the early stage, followed by a decline during the late stage (Supplementary Figure 14K). Indeed. Consistent with previous studies, SFRP1 was secreted by fibroblast cells, play a crucial role in driving the development of castration-resistant prostate cancer^1,2^.

**Two specific** **endothelial subsets promote** **prostate cancer progression**

Endothelial cells, intricately regulated by the TME, are pivotal in facilitating tumor growth and metastasis[38]. We further extracted and clustered the endothelial cells, identifying four distinct subtypes: Endothelial-c1-CLDN5, Endothelial-c2-ACKR1, Endothelial-c3-DCN, and Endothelial-c1-MKI67 (Supplementary Figure 15A, B, C). Using Slingshot and Monocle3, endothelial cells were projectedinto two branches, consisting of one root subtype (Endothelial-c4-MKI67), one intermediate subtype (Endothelial-c1-CLDN5), and two terminal subtypes (Endothelial-c2-ACKR1, Endothelial-c3-DCN) (Supplementary Figure 15D, E). Interestingly, both Endothelial-c2-ACKR1 and Endothelial-c3-DCN correlated with poor outcomes across multiple prostate cancer cohorts (Supplementary Figure 15J, K). DEGs analysis revealed that Endothelial-c2-ACKR1 highly expressed inflammatory factors and chemokines genes, such as CXCL1, CXCL10, IL6 (Supplementary Figure 15F). While Endothelial-c3-DCN highly expressed mesenchymal genes, such as COL1A1, SPARC, COL4A1 (Supplementary Figure 15F). In addition, GO enrichment analysis showed that Endothelial-c2-ACKR1 exhibited significant enrichment in myeloid cell homeostasis as well as leukocyte cell-cell adhesion (Supplementary Figure 15G). While endothelial-c3-DCN was enriched in collagen fibril organization, collagen metabolic process (Supplementary Figure 10G), with exhibiting high levels of both endothelial markers (PLVAP, PECAM1) and fibroblast markers (DCN, COL1A1, COL1A2) (Supplementary Figure 15B), suggesting endothelial-c3-DCN exhibited characteristics of both endothelial cells and fibroblasts. To explore the transitional dynamics between these endothelial cells and fibroblasts (Supplementary Figure 15H). Using Monocle2^3^, We found Endothelial-c3-DCN situated at an intermediate position along the pseudotime trajectory, bridging the transition between endothelial cells and two distinct fibroblast groups (Supplementary Figure 15I). The endothelial-to-mesenchymal transition (EndMT) process has been shown to play a pivotal role in orchestrating malignant tumor proliferation and facilitating metastasis^4^. Collectively, these findings suggest that endothelial-c2-ACKR1 promotes the progression of malignant cells by attracting peripheral blood cells, while endothelial-c3-DCN may contribute to a poor prognosis in patients with prostate cancer through the EndMT process.

**A special neutrophil subset** **possess immunosuppression phenotype in prostate cancer**

Based on the progressive increase of myeloid neutrophils along with lineage progression, we subdivided neutrophils into 11 subsets (Supplementary Figure 16A, B, C) and they were significantly enriched in GEMMs but not in WT (Supplementary Figure 16D). Among the seven neutrophil subsets, PMN-c6-IFIT1 showed a strong preference for distribution in PtRP (Supplementary Figure 16E) and was designated as tumor-related neutrophils (TANs) associated with significantly worse prognosis (Supplementary Figure 16F). Specifically, PMN exhibited a significant increase of CD274 expression, which encodes PD-L1, compared to other cell types (Supplementary Figure 16G), with PMN-c6-IFIT1 showing the highest expression (Supplementary Figure 16H). Gene ontology analysis of genes associated with PMN-c6-IFIT1 revealed enrichment in signaling pathways related to interferon-gamma (Supplementary Figure 16I). The expression of IFNG, which encodes the secreted protein interferon-gamma, showed a strongly positive correlation with CD274 in TCGA data (Supplementary Figure 16J). Additionally, IFNG was predominantly expressed by T/NK cells (Supplementary Figure 16K), suggesting that T/NK cells might secrete IFN-γ, contributing to the high PD-L1 expression in PMN-c6-IFIT1. Consistently, a study by Xue and colleagues recently demonstrated similar IFIT1+ neutrophil subsets showed high PDL1 expression can suppress T cell cytotoxicity liver tumors^5^. Based on these results, we propose that interferon-gamma secreted by lymphocytes could upregulate PD-L1 expression on TANs and shape an immunosuppressive microenvironment, promoting prostate cancer progression.
